# Type B and Type A influenza polymerases have evolved distinct binding interfaces to recruit the RNA polymerase II CTD

**DOI:** 10.1101/2022.02.04.479088

**Authors:** Tim Krischuns, Catherine Isel, Petra Drncova, Maria Lukarska, Alexander Pflug, Sylvain Paisant, Vincent Navratil, Stephen Cusack, Nadia Naffakh

## Abstract

During annual influenza epidemics, influenza B viruses (IBVs) co-circulate with influenza A viruses (IAVs), can become predominant and cause severe morbidity and mortality. Phylogenetic analyses suggest that IAVs (primarily avian viruses) and IBVs (primarily human viruses) have diverged over long time scales. Identifying their common and distinctive features is an effective approach to increase knowledge about the molecular details of influenza infection. The virus-encoded RNA-dependent RNA polymerases (FluPol_B_ and FluPol_A_) are PB1-PB2-PA heterotrimers that perform transcription and replication of the viral genome in the nucleus of infected cells. Initiation of viral mRNA synthesis requires a direct association of FluPol with the host RNA polymerase II (RNAP II), in particular the repetitive C-terminal domain (CTD) of the major RNAP II subunit, to enable “cap-snatching” whereby 5’-capped oligomers derived from nascent RNAP II transcripts are pirated to prime viral transcription. Here, we present the first high-resolution co-crystal structure of FluPol_B_ bound to a CTD mimicking peptide at a binding site crossing from PA to PB2. By performing structure-based mutagenesis of FluPol_B_ and FluPol_A_ followed by a systematic investigation of FluPol-CTD binding, FluPol activity and viral phenotype, we demonstrate that IBVs and IAVs have evolved distinct binding interfaces to recruit the RNAP II CTD, despite the CTD sequence being highly conserved across host species. We find that the PB2 627 subdomain, a major determinant of FluPol-host cell interactions and IAV host-range, is involved in CTD-binding for IBVs but not for IAVs, and we show that FluPol_B_ and FluPol_A_ bind to the host RNAP II independently of the CTD. Altogether, our results strongly suggest that the CTD-binding modes of IAV and IBV represent avian- and human-optimized binding modes, respectively, and that their divergent evolution was shaped by the broader interaction network between the FluPol and the host transcriptional machinery.

**Authors summary:** During seasonal influenza epidemics, influenza B viruses (IBVs) co-circulate with influenza A viruses (IAVs) and can cause severe outcomes. The influenza polymerase is a key drug target and it is therefore important to understand the common and distinctive molecular features of IBV and IAV polymerases. To achieve efficient transcription and replication in the nucleus of infected cells, influenza polymerases closely cooperate with the cellular RNA polymerase II (RNAP II) and interact with the repetitive C-terminal domain (CTD) of its major subunit. Here we gained new insights into the way IBV and IAV polymerases interact with the CTD of RNAP II. High-resolution structural data was used to perform structure-based mutagenesis of IBV and IAV polymerases followed by a systematic investigation of their interaction with RNAP II, transcription/replication activity and viral phenotype. Strikingly, we found that IBVs and IAVs have evolved distinct interfaces to interact with the host transcriptional machinery, in particular with the CTD of RNAP II. We provide evidence that these differences may have evolved as a consequence of the differences in IBV and IAV host range. Our findings are of significant importance with regard to the development of broad-spectrum antivirals that target the virus-host interface.

## Introduction

Influenza viruses are members of the *Orthomyxoviriridae* family and are classified into four genera: influenza A, B, C and D viruses. Influenza A viruses (IAVs) and influenza B viruses (IBVs) are of public health importance, as they co-circulate in humans with a seasonal epidemic pattern and cause a significant morbidity and mortality, especially in the aged or immunocompromised population [1]. IBV infections account for an estimated 23% of all influenza cases [2], can become predominant during annual influenza epidemics, and can cause severe disease in children [3]. IBVs have received less attention because, unlike IAVs which continuously circulate in a wide range of birds and mammalian species [4], they have no known potential to cause pandemics. Based on sequence analysis of the viral hemagglutinin, the evolutionary divergence between IBVs and IAVs was estimated to have occurred about 4000 years ago [5]. The recent identification of IBV-like viruses in non-mammalian vertebrate species suggest that IBVs and IAVs have actually diverged over much longer time scales [6]. IBVs and IAVs share the same genome organization of eight single-stranded negative RNA segments, and major features of the viral replication cycle such as transcription and replication of the viral genome in the nucleus of infected cells. However their genes have undergone functional divergence, as reflected notably by the lack of intertypic genetic reassortment [7]. To identify common and distinctive features of IBVs and IAVs is an effective approach to improve our understanding of the molecular mechanisms of influenza infection and our ability to fight influenza disease.

The genomic RNA segments of IAVs and IBVs are organized into viral ribonucleoprotein complexes (vRNPs) [8]. In the vRNP, the 5′ and 3′ terminal viral RNA sequences are associated with one copy of the RNA-dependent RNA polymerase complex (FluPol) while the RNA is covered by multiple copies of the viral nucleoprotein (NP) [9–11]. FluPol is a heterotrimer composed of PB1 (polymerase basic protein 1), PB2 (polymerase basic protein 2), and PA (polymerase acidic protein) [12], which replicates and transcribes the viral RNA in the nucleus of infected host cells. Replication is a primer-independent two-step process, which relies on *de novo* initiation by FluPol [13, 14]. In contrast, viral transcription is primer-dependent and results in the synthesis of 5′ capped and 3′ polyadenylated mRNAs, which are translated by the host translation machinery [15, 16]. Polyadenylation is achieved by stuttering of FluPol at a 5′ proximal oligo(U) stretch present on the genomic RNA [17, 18]. In contrast to other RNA virus polymerases, FluPol cannot synthesize 5′ cap structures [19]. In a process referred to as cap-snatching [20], FluPol binds the 5′ cap of nascent host RNA polymerase II (RNAP II) transcripts by the PB2 cap-binding domain. Then, the PA endonuclease domain [21] cleaves 10–15 nts downstream of the 5′ cap thereby generating primers that are used by FluPol to initiate transcription [18,19,22].

To perform cap-snatching, FluPol needs access to nascent capped RNAP II-derived RNAs, which represents a challenge as host cap structures are rapidly sequestered co-transcriptionally by the cap-binding complex [23]. The cellular RNAP II consists of 12 subunits [24], and the largest subunit (RPB1) is characterised by a unique long unstructured C-terminal domain (CTD) which in mammals consists of 52 repeats of the consensus sequence Tyr-Ser-Pro-Thr-Ser-Pro-Ser (Y_1_S_2_P_3_T_4_S_5_P_6_S_7_). Post-translational modifications of the CTD during the transcription process are controlling the spatiotemporal regulation of RNAP II transcription [25, 26]. FluPol binds specifically to S5 phosphorylated CTD (CTD pS5) [27, 28] and it was proposed that it targets RNAP II for cap-snatching in the paused elongation state, of which CTD pS5 is the hallmark modification [29–31].

Structural studies revealed bipartite CTD binding sites on the FluPol of influenza A, B and C viruses (FluPol_A_, FluPol_B_ and FluPol_c_) with notable differences from one type to another [32, 33]. However, the original crystal structure data for FluPol_B_ were of insufficient resolution and only one of the CTD binding sites could be modelled, therefore preventing functional studies. In this study, we report the first high-resolution co-crystal structure of FluPol_B_ bound to a CTD pS5 mimicking peptide that allows the modelling of both CTD-binding sites, one exclusively on PA also observed on FluPol_A_, and another, crossing from PA to PB2, specific for FluPol_B_. We used these novel data to perform structure-guided mutagenesis of FluPol_B_ and FluPol_A_, followed by a systematic investigation of cell-based CTD-binding, cell-based polymerase activity and plaque phenotype of recombinant viruses. Our findings demonstrate that type B and type A influenza polymerases have evolved distinct binding interfaces to recruit the RNAP II CTD, which is intriguing as the RNAPI II CTD is highly conserved across influenza host species. We find that the PB2 627 subdomain, a major determinant of FluPol-host cell interactions and IAV host-range, is involved in CTD-binding for IBVs but not for IAVs. Finally, we provide evidence for additional FluPol-RNAP II interactions that do not involve the CTD.

## Materials and Methods

### Purification, crystallisation, data collection and structure determination of FluPol_B_ with bound CTD peptide

Influenza B/Memphis/13/2003 polymerase, wild type or with the PA K135A mutation to eliminate endonuclease activity, was expressed and purified as described previously [22]. For crystals enabling high resolution visualisation of CTD binding in site 2B, FluPol_B_ PA mutant K135A at 9 mg ml^−1^ (35 µM) was mixed with 40 µM of nucleotides 1-13 vRNA 5’ end (5′-pAGUAGUAACAAGA-3′) and 1.8 mM 28-mer CTD peptide (YSPTpSPS)_4_ in a buffer containing 50 mM HEPES pH 7.5, 500 mM NaCl, 5% glycerol, 2 mM TCEP. Hanging drops for crystallisation were set up at 20 °C. Rod-shaped crystals growing up to 700 µm in length appeared one week after set-up in mother liquor containing 100 mM tri-sodium citrate and 13 % PEG 3350 with a drop ratio of 0.5 µl + 2 µl protein to well solution. Crystals were cryo-protected with additional 20 % glycerol and 1.8 mM CTD peptide in mother liquor and flash-frozen in liquid nitrogen. Data were collected on ESRF beamline ID29 and integrated with an ellipsoidal mask using AUTOPROC/STARANISO to an anisotropic resolution of 2.42-2.95 Å. The structure was solved using molecular replacement with PHASER [34] using PDB:5FMZ as model [35]. The model was iteratively corrected and refined using COOT [36] and REFMAC5 [37] and quality-controlled using MOLPROBITY [38]. See **Table 1** for data collection and refinement statistics.

**Table 1.** Crystallographic data collection and refinement statistics.

|  |  |  |  |
| --- | --- | --- | --- |
| **Crystal** | B/Memphis polymerase (WT) +3' vRNA 1-18 +5' vRNA 1-13 +13-mer capped RNA 28-mer CTD | B/Memphis polymerase (PA K135A) +5' vRNA 1-14 28-mer CTD | A/H7N9 polymerase core 201-PA_PB1_PB2-127 12-mer 5' vRNA pSer5 28-mer CTD peptide |
| **Diffraction Data** |  |  |  |
| Space group | $P3_221$ | $P2_1$ | $P2_1 2_1 2_1$ |
| Cell dimensions (Å) | a=b=200.82, c=257.02 | a=130.93, b=202.27, c=135.73, $\beta=110.54^\circ$ | a=76.47, b=144.13, c=336.20 |
| Wavelength (Å) | 0.966 | 1.254 | 0.968 |
| Beamline (ESRF) | ID30A1 | ID29 | ID29 |
| No. Crystals | 1 | 1 | 1 |
| Resolution range (last shell) (Å) | Isotropic 3.56 Å<br>Ellipsoidal 3.12 Å<br>76.85.0-3.12 (3.32-3.12) | Isotropic 2.95 Å<br>Ellipsoidal 2.42 Å<br>127.1-2.42 (2.69-2.42) | Ellipsoidal 3.41 Å<br>49.21-3.41 (3.57-3.41) |
| Completeness (last shell) (%) | Overall 73.1<br>Ellipsoidal: 91.6 (67.9) | Overall 61.4<br>Ellipsoidal: 89.1 (64.0) | Overall 73.3<br>Ellipsoidal 94.8 (88.2) |
| R-sym (last shell) | 0.128 (1.30) | 0.112 (0.618) | 0.283 (1.873) |
| R-meas (last shell) | 0.149 (1.49) | 0.136 (0.769) | 0.306 (1.946) |
| I/σI (last shell) | 8.0 (1.7) | 7.7 (1.9) | 7.7 (1.5) |
| CC(1/2) (last shell) | 0.994 (0.536) | 0.992 (0.630) | 0.995 (0.696) |
| Redundancy (last shell) | 3.9 (3.9) | 3.0 (2.5) | 14.1 (14.6) |
| **Refinement** |  |  |  |
| Reflections (total)<br>For refinement: work (free) | 77848<br>73897 (3951) | 154519<br>146946 (7573) | 37875<br>36023 (1852) |
| R-work (last shell) | 0.240 (0.369) | 0.223 (0.328) | 0.220 (0.313) |
| R-free (last shell) | 0.273 (0.397) | 0.257 (0.257) | 0.273 (0.358) |
| No of non-hydrogen atoms | 18481 | 35955 | 19969 |
| **Validation** |  |  |  |
| RMS(bonds) | 0.002 | 0.003 | 0.003 |
| RMS(angles) | 1.170 | 1.198 | 1.207 |
| Average B-factor (Å<sup>2</sup>) | 109.0 | 57.89 | 86.16 |
| Ramachandran favored (%) | 92.59 | 96.18 | 92.65 |
| Ramachandran outliers (%) | 0.32 | 0.09 | 0.79 |
| Molprobrity score | 1.50 | 1.76 | 1.96 |
| Clash score | 1.20 | 2.74 | 2.83 |

For crystals enabling simultaneous visualisation of CTD binding in sites 2A and 2B, wild type FluPol_B_ at 11.7 mg ml^−1^ (45 µM) was mixed with 52 µM of nucleotides 1-14 (5′-pAGUAGUAACAAGAG-3′) and 1-18 (5′-UAUACCUCUGCUUCUGCU-3′) of respectively the vRNA 5’ and 3’ ends, 104 µM of capped 13-mer (5’-m^7^GpppAAUCUAUAAUAGC-3’), and 380 µM CTD 28-mer Ser5P peptide, in buffer containing 50 mM HEPES pH 7.5, 500 mM NaCl, 5 % glycerol, 2 mM TCEP. Hanging drops for crystallisation were set up at 4° C. Diamond-shaped crystals growing up to 200 µm in size appeared in two to three weeks in mother liquor containing 200 mM di-ammonium phosphate, and 100 mM sodium acetate at pH 4.4, with a drop ratio of 1 µl + 2 µl protein to well solution. The drops were soaked with 840 µM CTD peptide for 17 days. Crystals were cryo-protected with an additional 30 % glycerol and 885 µM peptide in mother liquor and flash-frozen in liquid nitrogen. Data were collected on ESRF beamline ID30A1 (MASSIF) and processed and refined as described above, using PDB:5MSG as model for molecular replacement. See **Table 1**.

### Structure determination of FluPol_A_(H7N9) core with bound CTD peptide

The core of influenza A/Zhejiang/DTID-ZJU01/2013(H7N9) polymerase comprising PA 201-716, PB1 full-length, PB2 1-127 was expressed and purified from insect cells as described previously [18]. A/H7N9 polymerase core at a concentration of 9 mg/ml was co-crystallised with 60 µm of a 12-mer of the vRNA 5’ end (5’-pAGUAGUAACAAG) in sitting drops at 4° C in conditions of 0.1 M Tris pH 7.0, 13 % PEG 8K, 0.2 M MgCl_2_, 0.1 M guanidine hydrochloride with drop mixing ratios of 1:2 (protein:well). Crystals grew typically within 4-5 days and diffracted to around 3.5 Å resolution. A four-repeat pS5 CTD mimicking peptide (Tyr-Ser-Pro-Thr-pSer-Pro-Ser)_4_ was soaked into existing crystals at a concentration of ∼2 mM over a period of 24 h. Data were collected on ESRF beamline ID29 and processed and refined as described above, using previously described apo-H7N9 core structure ([39], PDB:6TU5) as model for molecular replacement. See **Table 1** for data collection and refinement statistics.

### Cells

HEK-293T cells (purchased at ATCC, CRL-3216) were grown in complete Dulbecco’s modified Eagle’s medium (DMEM, Gibco) supplemented with 10% fetal bovine serum (FBS) and 1% penicillin-streptomycin. MDCK cells (provided by the National Influenza Center, Paris, France) were grown in Modified Eagle’s medium (MEM, Gibco) supplemented with 5% FBS and 1% penicillin-streptomycin.

### Plasmids

The reverse genetics plasmids derived from the IAV A/WSN/33 (WSN) [40] and the IBV B/Brisbane/60/2008 [41] were kindly provided by Pr. G. Brownlee (Sir William Dunn School of Pathology, Oxford, UK) and Pr. D. Perez (College of Veterinary Medicine, University of Georgia), respectively. For polymerase activity assays, a pPolI-Firefly plasmid encoding the Firefly luciferase sequence in negative polarity flanked by the 5’ and 3’ non-coding regions of either the IAV or IBV NS segment was used. The pRenilla-TK plasmid (Promega) was used as an internal control. The WSN pcDNA3.1-PB2, -PB1, -PA plasmids [32] and B/Memphis/13/2003 (Memphis) pcDNA3.1-PB2, -PB1, -PA and -NP plasmids [42] were described previously. The WSN-NP Open Reading Frame (ORF) was subcloned into the pCI-plasmid. The WSN and Memphis pCI-PB2-G1 and pCI-PA-G1 plasmids used for *Gaussia* luciferase complementation assays were constructed as described previously [43]. The RPB1 ORF was obtained from the Human ORFeome resource (hORFeome v3.1), fused to G2 at the N-terminus by PCR and subcloned into pcDNA3.1 (G2-RPB1). The CTD repeats 4 to 51 were deleted from the G2-RPB1 construct by PCR (G2-RPB1ΔCTD). The pCI-RPB2-G2 construct was kindly provided by Dr. B. Delmas (INRAE, Jouy-en-Josas). The wild-type full-length CTD sequences was fused at the C-terminus to an SV40 NLS by PCR using the G2-RPB1 plasmid as a template. The resulting amplicon was cloned in frame downstream the G2 sequence into the pCI vector (G2-CTD). A sequence in which each CTD serine 5 residue was replaced by an alanine was ordered as synthetic gene (GenScript) and subcloned in place of the wild-type CTD sequence into the G2-CTD construct (G2-CTD-S5A). The pCI-G2-NUP62 plasmid was described previously [44]. Mutations were introduced by an adapted QuickChange™ site-directed mutagenesis (Agilent Technologies) protocol [45]. Primers and plasmid sequences are available upon request.

### Protein complementation and minigenome assays

HEK-293T cells were seeded in 96-well white opaque plates (Greiner Bio-One) the day before transfection. For the split-luciferase complementation assays, cells were co-transfected in technical triplicates with 25 ng plasmid encoding the polymerase subunits PB2, PB1 and PA (either PB2-G1 or PA-G1, respectively) and 100 ng of the G2-tagged targets (CTD, RPB1 or RPB2, respectively) using polyethyleneimine (PEI-max, #24765-1 Polysciences Inc). When indicated, the CDK7 inhibitor BS-181-HC (Tocris Bioscience) was added 24 hours post-transfection (hpt) at a final concentration of 20 µM for 1 h. DMSO 0.2 % was used as a control. Cells were lysed 20-24 hpt in *Renilla* lysis buffer (Promega) for 45 min at room temperature under steady shaking (650 rpm) and the *Gaussia princeps* luciferase enzymatic activity was measured on a Centro XS LB960 microplate luminometer (Berthold Technologies, reading time 10 s after injection of 50 µl Renilla luciferase reagent (Promega). For the minigenome assays, cells were co-transfected in technical triplicates with 25 ng of each pcDNA3.1 PB2, PB1, PA, in conjunction with 50, 10 and 5 ng of the pCI-NP, pPolI-Firefly and pTK-Renilla plasmids, respectively. Luciferase activities were measured 20-24 hpt using the the Dual-Glo Luciferase Assay system (Promega) according to the manufacturer’s instructions.

### Antibodies and immunoblots

Total cell lysates were prepared in RIPA cell lysis buffer as described before [46]. Immunoblot membranes were incubated with primary antibodies directed against CTD-pS5 (Active Motif, 3EB), CTD-pS2 (Active Motif, 3E10), *Gaussia princeps* luciferase (New England Biolabs, #E8023) or Tubulin (Sigma-Aldrich, B-5-1-2), and subsequently with the according HRP-tagged secondary antibodies (Jackson Immunoresearch). Membranes were revealed with the ECL2 substrate according to the manufacturer’s instructions (Pierce) and chemiluminescence signals were acquired using the ChemiDoc imaging system (Bio-Rad) and analysed with ImageLab (Bio-Rad).

### Production and characterisation of recombinant viruses

The recombinant viruses were produced by transfection of a co-culture of HEK-293T and MDCK cells as described previously [40, 41]. The reverse genetics supernatants were titrated on MDCK cells in a standard plaque assay as described before [47]. Plaque diameters were measured upon staining with crystal violet using Fiji [48].

### *In vitro* endonuclease and transcription activity assays

RNA for the activity assays was produced *in vitro* with T7 polymerase. Recombinant polymerases used corresponding to A/little yellow-shouldered bat/Guatemala/060/2010 and B/Memphis/13/2003 were purified as previously described [42]. 23 nt RNA (5’-GAAUCUAUACAUAAAGACCAGGC-3’) was capped with vaccinia capping enzyme and 2’-O-methyltransferase (NEB) and radiolabelled with [α ^32^P]-GTP. For the endonuclease assay 25 nM FluPol_A_ and 50 nM FluPol_B_ were incubated with 1.2-fold molar excess of v5’ RNA (FluPol_A_ : 5’-pAGUAGAAACAAGGC-3’, FluPol_B_ : 5’-pAGUAGUAACAAGAG-3’) and the capped RNA in reaction buffer containing 50 mM HEPES pH 7.5, 150 mM NaCl, 5 mM MgCl_2_, 2 mM tris(2-carboxyethyl)phosphine (TCEP). Transcription reactions were performed with 50 nM FluPol_A_ or FluPol_B_ in the reaction buffer, supplemented with 150 nM v3’ template RNA (FluPol_A_: 5’-AGUUUGCCUGCUUCUGCU-3’, FluPol_B_: 5’-UAUACCUCUGCUUCUGCU-3’) and 250 µM NTP mix (ThermoFisher). 50 µM CTD peptides were added at concentrations corresponding to at least a 10-fold excess over the K_D_ of the lowest measured affinity for a two-repeat peptide. Two- and four-CTD repeat peptides were purchased from Covalab and six-repeat CTD peptide was synthesised at the Chemical Biology Core Facility at EMBL Heidelberg. Reactions were incubated at 30 °C for 30 min and quenched with RNA loading dye (formamide, 8 M urea, 0.1 % SDS, 0.01 % bromophenol blue (BPB), 0.01 % xylene cyanol (XC)), supplemented with 50 mM EDTA and boiled at 95 °C. The reaction products were separated on 20 % denaturing acrylamide gel (containing 8 M urea) in Tris-Borate-EDTA (TBE) buffer, exposed on a Storage Phosphor screen and recorded with a Typhoon reader. DECADE_TM_ marker was used as ladder.

### CTD sequences and alignment

The RPB1 CTD domain amino acid sequences of *Homo sapiens* (NP_000928.1), *Sus scrofa* (XP_020923484.1), *Equus caballus* (XP_014584045.2), *Canis lupus* (XP_038521325.1), and *Mus musculus* (NP_001277997.1) host species were obtained from the RefSeq database [49]. As the predicted RefSeq sequences available for the *Gallus gallus* (XP_040551262) and *Anus platyrhyncos* (XM_038172734) RPB1 subunits were only partial, we designed a targeted protein sequence assembly strategy data based on RNA-seq and/or WGS SRA public data available for these two species. To obtain the *Gallus gallus* RPB1 complete sequence (1969 aa), we first aligned Illumina RNA-seq short reads (ERR2664216) on the human RefSeq curated protein sequence (NP_000928) using DIAMOND algorithm [50], and then used the aligned reads for subsequent Trinity transcript assembly (“–longreads XP_040551262” option to use the partial sequence as a guide) followed by Transdecoder for the ORF prediction [51]. The *Anus platyrhyncos* RPB1 complete sequence (1970 aa) was obtained by aligning Illumina RNA-seq short reads (SRR10176883) and PACBIO long reads (SRR8718129, SRR8718130) on the JACEUL010000271.1 genomic scaffold by using respectively HISAT2 [52] and minimap2 [53] followed by Stringtie2 [54] with the “–mix” option to allow hybrid de novo gene assembly. The CTD sequences were aligned with SnapGene ® 6.0 and visualised by Espript 3.0 [55].

## Results

### Cocrystal structures reveal distinct CTD binding sites in FluPol_B_ and FluPol_A_

Previous structural studies using a four repeat CTD pS5 peptide mimic (YSPTpSPS)_4_ [32] revealed two distinct CTD binding sites on FluPol_B_, denoted site 1B and site 2B. Site 1B, exclusively on the PA subunit and in which the pS5 phosphate is bound by PA basic residues K631 and R634, is essentially the same as site 1A for bat influenza A polymerase [32] and is thereafter named site 1AB. Site 2B, which extends across the PA-PB2 interface, is unique to FluPol_B_ and distinct from site 2A for FluPol_A_, which is again exclusive to the PA subunit [32]. However, the original crystal structure data for FluPol_B_ were of insufficient resolution to be able to construct a model for the CTD peptide in site 2B, nor even to define its directionality. To overcome this limitation, we co-crystallised the four repeat pS5 peptide with influenza B/Memphis/13/2003 polymerase in a different *P*2_1_ crystal form, previously used to obtain a structure with the 5’ end of the vRNA [35], and measured anisotropic diffraction data to a resolution of 2.42-2.95 Å (**Table 1**). The resultant map, which contains two heterotrimers in the asymmetric unit, showed clear electron density in site 2B for both trimers (**Figure S1A**), into which an unambiguous model for the CTD peptide could be built (**Figure 1A and Figure S2A**). Only very weak density for the CTD peptide is observed in site 1AB, perhaps because of competition with a phosphate bound at the position of the phosphoserine. To reconfirm that sites 1B and 2B could be occupied simultaneously, we re-crystallised full promoter-bound FluPol_B_ with the CTD peptide in the original P3_2_21 crystal form, but this time with a capped primer and at lower pH. Under these conditions, the extremity of the vRNA 3’ end is in the RNA synthesis active site [56]. Anisotropic diffraction data to a resolution of 3.12-3.56 Å was measured and the resultant map showed clear electron density for the CTD peptide bound in both sites 1AB and 2B **(Figure 1B and Table 1)**, as reported previously for this crystal form [32] but with slightly improved resolution. Unexpectedly, the CTD peptides bound in site 1AB and site 2B are orientated such that they cannot be linked by the shortest path, as this would be between both N-termini, which are ∼ 17 Å apart, whereas the straight-line distance between the C-ter of site 1AB and N-ter of site 2B is ∼ 36 (44) Å. These distances suggest that a minimum of 6, probably 7, heptad repeats would be required to occupy both sites contiguously (**Figure 1B,** dotted red line). This contrasts with the situation in FluPol_A_, where the peptide directionality in sites 1AB and 2A allow them to be linked by the shortest path, implying that four heptad repeats is sufficient to occupy both sites (**Figure 1C**) [32].

**Figure 1.**
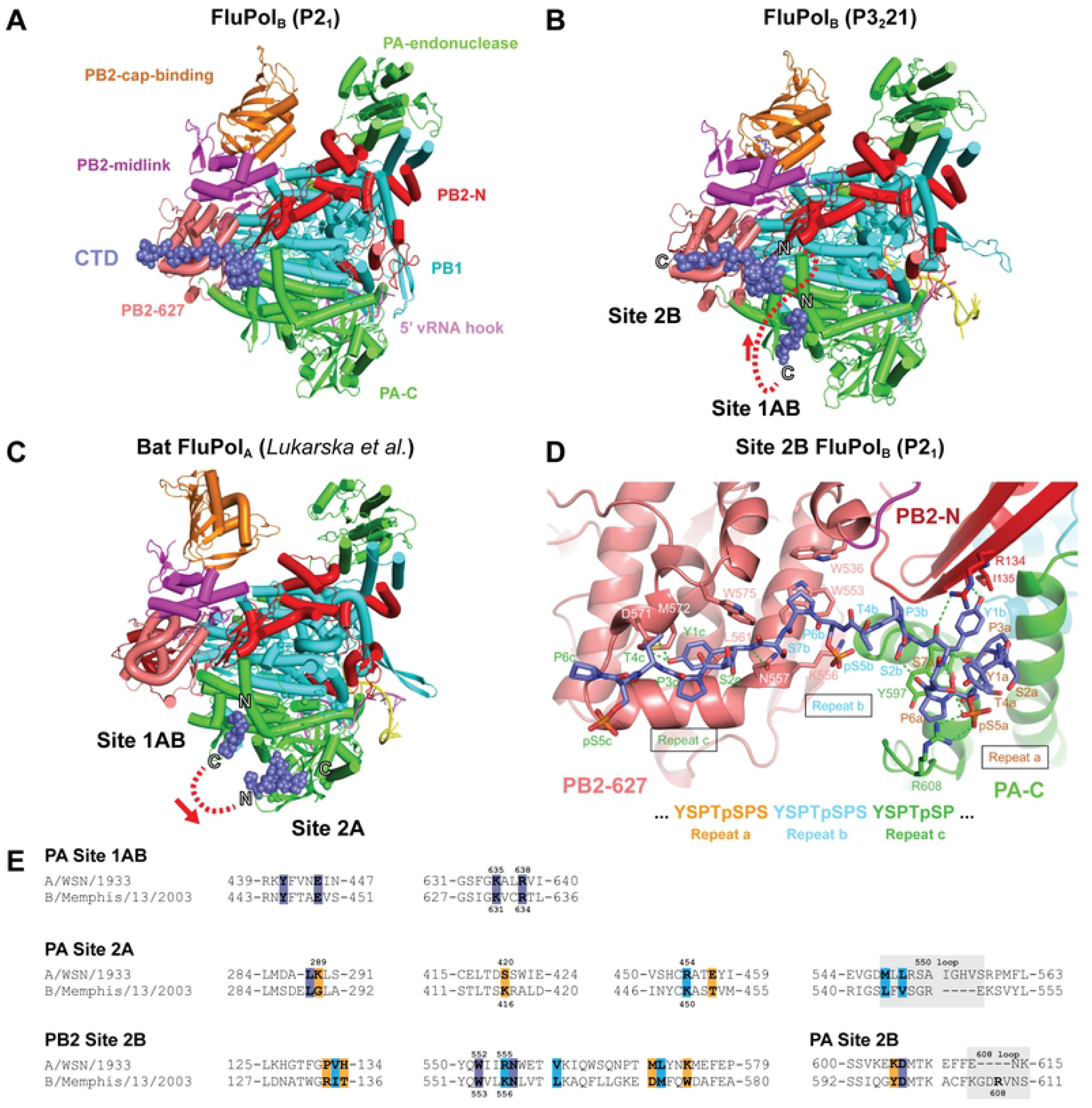
Structural analysis of CTD binding to influenza B polymerase. **A.** Overall view of the crystal structure of influenza B polymerase, with bound vRNA 5’ hook (pink) and CTD peptide mimic (slate blue spheres) in site 2B. Ribbon diagram of the polymerase with PA (green), PB1 (cyan), PB2-N (red), PB2-cap-binding (orange), PB2-midlink (magenta), PB2-627 (deep salmon). **B.** Overall view of the crystal structure of influenza B polymerase with bound promoter (pink and yellow), capped primer (blue) and CTD peptide mimic (slate blue spheres) bound in sites 1AB and 2B. The polymerase is coloured as in (A). The N and C-termini of the two CTD fragments are marked and the red dotted line shows the shortest connection between them with directionality indicated by the arrow. **C.** Overall view of the crystal structure of bat influenza A polymerase with bound promoter and CTD peptide mimic (slate blue spheres) bound in sites 1AB and 2A ([32], PDB: 5M3H). The colour code is as in (A). The N and C-termini of the two CTD fragments are marked and the red dotted line shows the shortest connection between them with directionality indicated by the arrow. **D.** Details of the interaction between key residues of the influenza B polymerase PA subunit (green), PB2-N (red) and PB2-627 (deep-salmon) with the CTD peptide (slate blue sticks) in site 2B. Three CTD repeats denoted a (orange), b (cyan) and c (dark green) are involved in this interaction. Hydrogen bonds are indicated as dotted green lines. **E.** Sequence alignment of the CTD binding sites in the A/WSN/33 (A0A2Z5U3X0) and B/Memphis/13/2003 (Q5V8X3) polymerase subunits PA and PB2. Protein sequences were obtained from UniProt (https://www.uniprot.org/) and aligned with SnapGene ® 6.0. Key residues for CTD binding are indicated in bold. Identical, similar and non-similar residues are highlighted in dark blue, light blue and orange, respectively. Residue submitted to mutagenesis in this study are indicated with their numbers above (FluPol_A_) and below (FluPol_B_) the alignment, respectively.

Three repeats (designated repeats a, b and c) of the CTD peptide (i.e. Y1aS2aP3aT4a**pS5**aP6aS7a-Y1bS2bP3bT4b**pS5**bP6bS7b-Y1cS2cP3cT4cpS5cP6c) are visible in site 2B in both structures, including two well-defined phosphoserines (in bold). The N-terminal part of the CTD peptide (Y1a-S2b) forms a compact structure comprising two successive proline turns stabilised by four intra-peptide hydrogen bonds, with P3a stacked on Y1b and P6a stacked on PA/Y597 (**Figure 1D)**. PB2 R132 partially stacks against the other side of the Y1b sidechain, whose hydroxyl group hydrogen bonds to the main-chain of PB2 I135. The phosphate of pS5a forms a strong salt-bridge with PA R608 as well as hydrogen bonding with S7a. FluPol_B_-specific PA R608 is in a four-residue insertion (606-GDRV-609) compared to FluPol_A_, with hydrogen bond interactions from PA D607 and N611 positioning the side-chain of PA Y597 under the CTD peptide. This configuration of residues seems specifically designed to accommodate the compactly folded CTD peptide. Interestingly, recently identified FluPol_B_-like polymerases from fish and amphibians [57] also possess the four-residue insertion in PA. However, only in the Wuhan spiny-eel influenza virus polymerase, which is remarkably similar to human FluPol_B_ polymerase, are all the functional residues Y597, D607, R608, N611 conserved [57, 58]. The rest of the CTD peptide (P3b-T4c) has an extended conformation and lies across the PB2 627-domain (**Figure 1D**). To create the CTD binding surface requires concerted side chain reorientations of PB2 W553, M572 and W575 (**Figure 1D and Figure S2A**), allowing P6b to pack on W553 and Y1c on M572 and L561, with its hydroxyl group hydrogen bonding to D571. PB2 K556 forms a salt bridge with pS5b.

Most functional studies on CTD are performed with human or avian influenza A polymerase, whereas CTD binding has only been structurally characterised for bat A/little yellow-shouldered bat/Guatemala/060/2010(H17N10) polymerase [32] and C/Johannesburg/1/1966 [33]. Although sequence alignments and mutational studies strongly suggest that the mode of CTD binding is conserved for all IAV-like polymerase [32], we attempted to confirm this by determining the structure of a CTD mimicking peptide bound to influenza A/Zhejiang/DTID-ZJU01/2013(H7N9) polymerase. Previously, we have reported crystals of the A/H7N9 core (PA 201-716, PB1 full-length, PB2 1-127) in the apo-state, which forms symmetrical dimers as described elsewhere [18,59,60]. Here, we soaked the four-repeat pS5 CTD peptide mimic into co-crystals of H7N9 core with the vRNA 5’ hook. The crystals diffracted to a maximum resolution of 3.41 Å (**Table 1**) and again contain symmetrical dimers of the polymerase core. We observed clear electron density, not only for the 5’ hook, but also for the CTD peptide bound in site 2A (**Figure S1B and Figure S2B**), essentially identically bound as previously seen for bat influenza A/H17N10 polymerase (**Figure S2B**). However, there was no CTD peptide bound in site 1AB. The most likely explanation for this is that in the symmetrical dimeric form of influenza A (core only, or full trimer), both polymerases are in the so-called ‘dislocated’ conformation [18] with an open active site. In particular, PA regions 425-452 and 586-618 are rotated by ∼20°, compared to the active, monomeric promoter bound state (e.g. A/H3N2 polymerase structure, [59], PDB:6RR7). This particularly affects the position of key site 1AB binding site residues Y445, E449 and F612 (**Figure S2C**), thus preventing CTD at this site, while not affecting binding to site 2A.

A sequence alignment of the CTD binding sites 1AB, 2A, and 2B from the representative influenza A and B viruses used hereinafter is shown in **Figure 1E** (sequence alignments of the full-length PB2, PA and PB1 subunit are provided in **Figure S3**). Key residues for CTD binding are mostly conserved (dark and light blue) in site 1AB, non-conserved in site 2A and partially conserved in site 2B.

### The FluPol-CTD interaction can be monitored using a cell-based luciferase complementation assay

To confirm the structural findings of this study and investigate the distinctive features of CTD binding sites in FluPol_B_ and FluPol_A_ in the cellular context, we set up a CTD-binding assay using the *Gaussia princeps* luciferase trans-complementing fragments (G1 and G2) [43]. The full-length CTD was fused to G2 and the SV40 nuclear localization signal (G2-CTD). PB2 or PA were fused to G1 at their C-terminus (PB2-G1 and PA-G1, respectively, schematically represented in red in **Figure 2A** and **2B**) as FluPol was shown to retain activity when tagged at these sites [44]. Upon co-expression of G2-CTD and the three polymerase subunits (including PB2-G1 or PA-G1), a luminescence signal resulting from the FluPol-CTD interaction was measured, which was generally higher for FluPol_B_ compared to FluPol_A_ (**Figure 2A** and **2B**). The interaction signal decreased when PB1 was omitted, in agreement with previous reports that the FluPol-CTD interaction depends on FluPol assembly [28], and it was independent of FluPol catalytic activity (**Figure 2A** and **2B**, PB1 D444A-D445A mutant [61]). When key CTD-contacting residues of FluPol_A_ were mutated (PA K289A and R638A [32]), the interaction signal was significantly decreased compared to PA wt (**Figure 2C** and **2D**). To test whether the FluPol-CTD binding assay reflects the dependency on the phosphorylation of the CTD S5 moiety, all S5 residues of the CTD were mutated to alanine (schematically represented in **Figure 2E**), which prevented S5 phosphorylation as documented by western blot (**Figure 2F** and **G, bottom**). Although the wt and S5A CTD showed similar steady-state levels of expression, the binding of the S5A CTD to FluPol_B_ and FluPol_A_ was significantly decreased compared to WT CTD. Consistently, pharmacological inhibition of CDK7, which represents the major kinase for CTD S5 phosphorylation [62], specifically reduced FluPol_A/B_ binding to the CTD but not to the FluPol_A_ interaction partner NUP62 [44] (**Figure S4A and S4B**). Overall, these data demonstrate that the FluPol-CTD interaction can be accurately monitored in cells using our split-luciferase complementation assay conditions.

**Figure 2.**
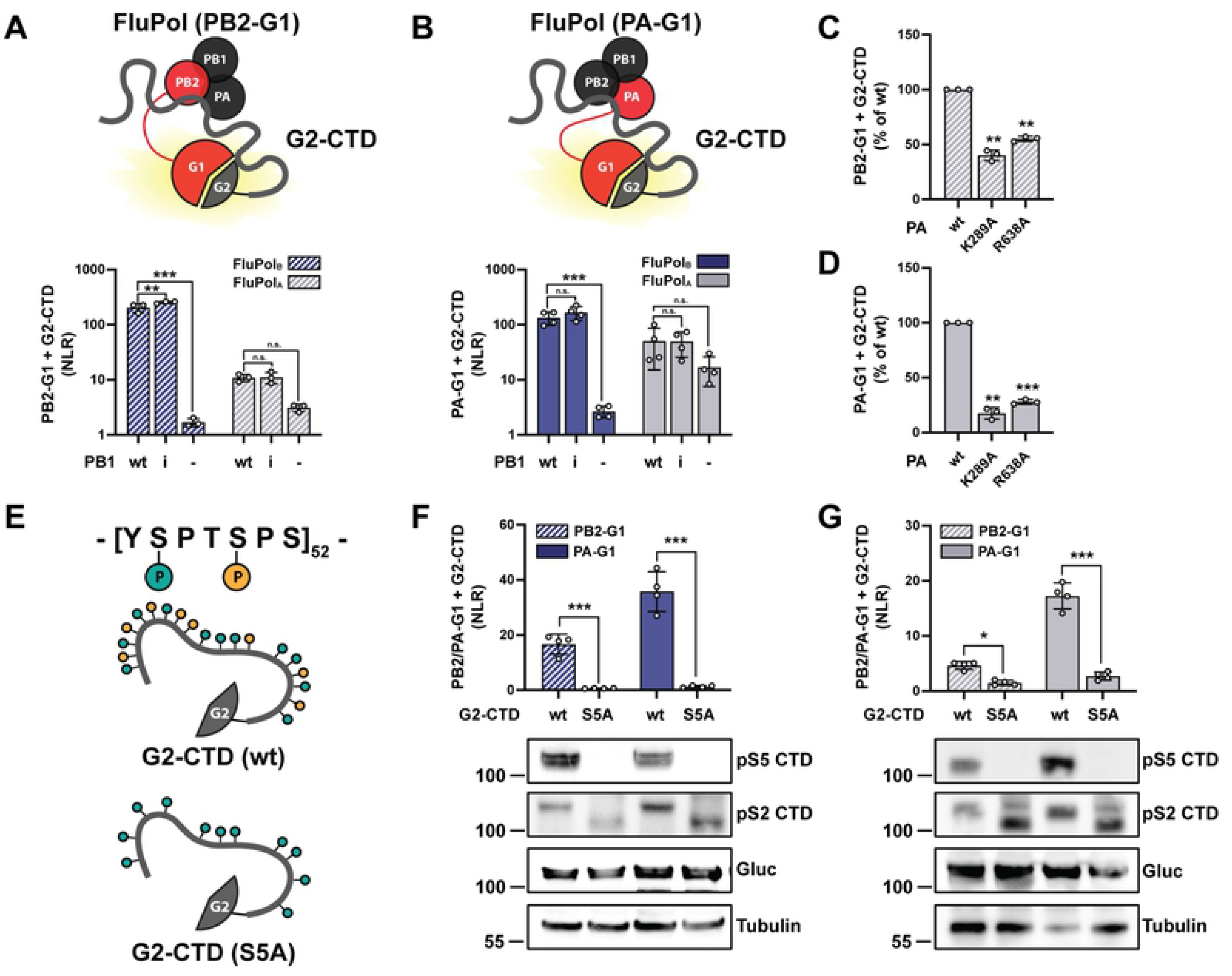
*Gaussia princeps* luciferase-based FluPol - CTD binding assay. **A-B.** G2-CTD was expressed by transient transfection in HEK-293T cells together with PB2, PB1 and PA of FluPol_B_ (B/Memphis/13/2003, blue bars) or FluPol_A_ (A/WSN/33, grey bars). Either PB2 (A, hatched bars) or PA (B, filled bars) were C-terminally tagged in frame with G1. As controls, the wild-type (wt) PB1 was replaced by the catalytic inactive PB1 D444A D445A mutant (i) or was omitted (-). Luciferase activities were measured in cell lysates at 24 hpt. Normalised luciferase ratios (NLRs) were calculated as described in the Materials and Methods section. The data shown are the mean ± SD of at least three independent experiments performed in technical triplicates. **p ≤ 0.002; ***p ≤ 0.001 (two-way ANOVA; Dunnett’s multiple comparisons test). **C-D.** The CTD binding of FluPol_A_ mutants PA K289A and R638A was investigated. HEK- 293T cells were transfected as described in (A) and (B), respectively. Relative Light Units (RLUs) are expressed as percentages relative to the FluPol_A_ PA wt. The data shown are the mean ± SD of three independent experiments performed in technical triplicates. **p ≤ 0.002; ***p ≤ 0.001 (one-way ANOVA; Dunnett’s multiple comparisons test). **E.** Schematic representation of the CTD constructs used in (F) and (G): the wild-type G2-CTD (wt, top) and the G2-CTD in which all serine 5 residues were replaced with an alanine (S5A, bottom). **F-G.** The interaction of the wt or the S5A mutated CTD to FluPol_B_ (F) or FluPol_A_ (G) was investigated by transient transfection in HEK-293T cells as described in (A). The data shown are the mean ± SD of four independent experiments performed in technical triplicates. *p ≤ 0.033; ***p ≤ 0.001 (two-way ANOVA; Sidak’s multiple comparisons test). In parallel, cell lysates were analysed by western blot using antibodies specific for the pS5 or pS2 CTD, *G.princeps* luciferase (Gluc) and tubulin.

### Structure-driven mutagenesis confirms FluPol_B_ and FluPol_A_ have distinct CTD binding modes on PA

To systematically assess *in vivo* FluPol-CTD binding, we mutated key residues forming the CTD binding sites in the FluPol_B_ and/or FluPol_A_ co-crystal structures and measured the impact of these mutations on CTD-binding using the split-luciferase complementation assay described above (**Figure 2**). In parallel, we investigated polymerase activity in a minireplicon assay, and we rescued recombinant mutant IBVs and IAVs and measured plaque diameters on reverse genetic supernatants as a read-out for viral growth capacity. The FluPol_A_ residue nature and numbering is used in the text and figures, except when indicated.

The structure and key residues of the CTD binding site 1AB are conserved between FluPol_B_ and FluPol_A_ (**Figure 1E** and **Figure 3A**). We mutated the pS5 interacting residues PA K635 and R638 to alanines. The mutations did not affect PA accumulation levels (**Figure 3B**) but significantly decreased *in vivo* binding to the full-length CTD for both FluPol_B_ and FluPol_A_ (**Figure 3C and Figure S5A**), which is in line with biochemical data obtained *in vitro* with CTD-mimicking peptides [32]. Consistently, the corresponding recombinant mutant IBVs and IAVs were attenuated or could not be rescued (**Figure 3D**). Noteworthy, the impact of site 1AB mutations on the viral phenotype and FluPol activity was generally lower for IBV than IAV mutants (**Figure 3D** and **3E**) although the defect in CTD binding was more pronounced for FluPol_B_ (**Figure 3C**), suggesting that FluPol_B_ activity is less reliant on CTD binding at site 1AB.

**Figure 3.**
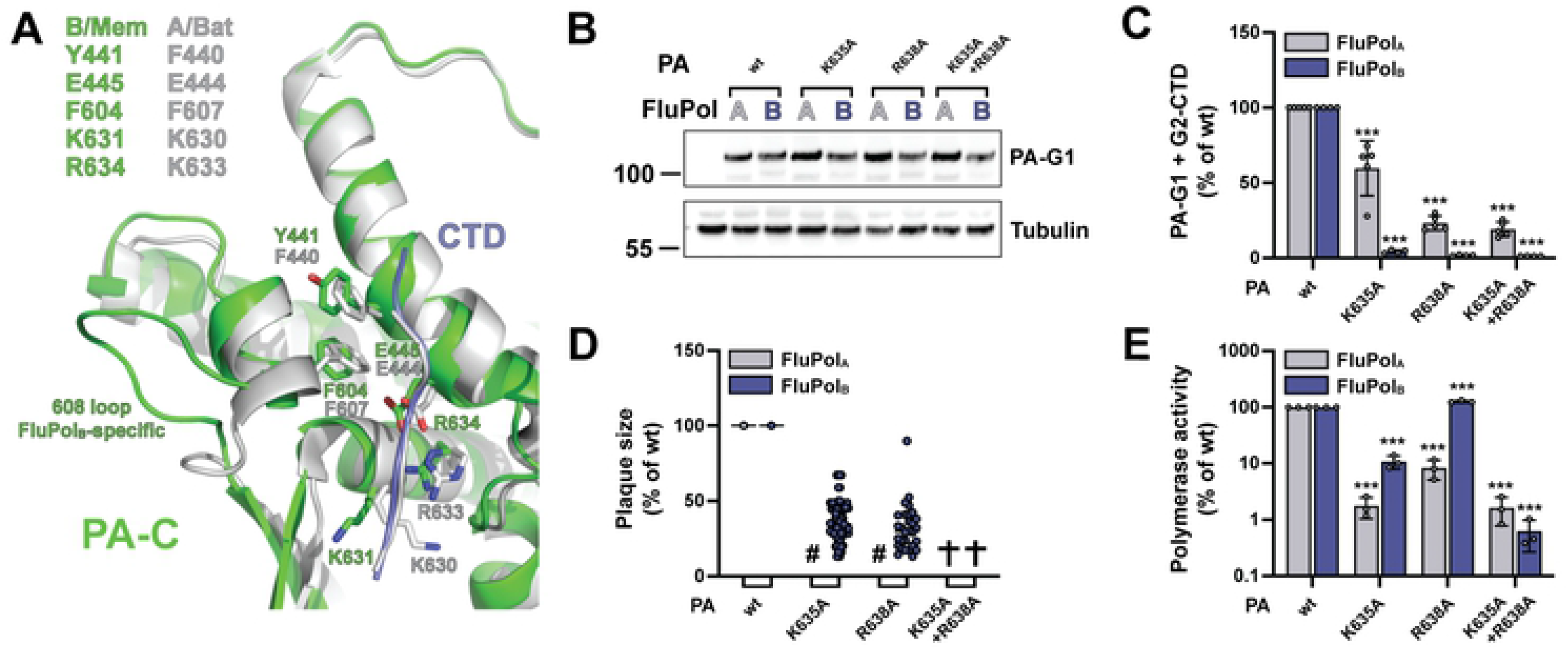
FluPol_B_ and FluPol_A_ CTD-binding mode at site 1AB. **A.** Superposition of the similar CTD binding in sites 1AB on the PA subunit for influenza B (B/Memphis/13/2003, green) and bat influenza A (A/little yellow-shouldered bat/Guatemala/060/2010(H17N10), light grey) polymerases with the CTD peptide as a thin tube (respectively slate blue and light grey). Key conserved residues are indicated in their respective colours, as well as the FluPol_B_-specific insertion (PA 608 loop) that is important for part of site 2B. See sequence alignment in Figure 1E. **B.** HEK-293T cells were transfected with the indicated FluPol_A_ (A/WSN/33) and FluPol_B_ (B/Memphis/13/2003) site 1AB mutants, which were C-terminally tagged with the G1 fragment. Cells were lysed at 24 hpt and analysed by western blot using antibodies specific for *G.princeps* luciferase (PA-G1) and tubulin. The residue numbering corresponds to FluPol_A_ (A/WSN/33). **C.** *In vivo* CTD binding of the indicated mutants of FluPol_A_ (A/WSN/33, grey bars) and FluPol_B_ (B/Memphis/13/2003, blue bars). The G2-tagged CTD was expressed by transient transfection in HEK-293T cells together with PB2, PB1 and PA-G1. RLUs are expressed as percentages relative to wt FluPol_A/B_. The data shown are mean ± SD of three independent experiments performed in technical triplicates. ***p ≤ 0.001 (two-way ANOVA; Dunnett’s multiple comparisons test). **D.** Characterisation of recombinant IAV (A/WSN/33, grey dots) and IBV (B/Brisbane/60/2008, blue dots) viruses. Recombinant viruses with the indicated mutations were generated by reverse genetics as described in the Material and Methods section. Reverse genetic supernatants were titrated on MDCK cells, stained at 72 hpi and plaque diameters were determined using the Fiji software. Each dot represents the diameter of a viral plaque relative to the mean plaque size of IAV wt or IBV wt recombinant virus. (#) not measurable pinhead-sized plaque diameter; (✞) no viral rescue. **E.** Polymerase activity of CTD-binding site 1AB mutants. FluPol_A_ (A/WSN/33, grey bars) or FluPol_B_ (B/Memphis/13/2003, blue bars) was reconstituted in HEK-293T cells by transient transfection of PB2, PB1, PA, NP and a model RNA encoding the Firefly luciferase flanked by the 5’ and 3’ non-coding regions of the IAV or IBV NS segments, respectively. As an internal control, a RNA-Polymerase II promotor driven Renilla plasmid was used. Luminescence was measured at 24 hpt as described in the Material and Methods section. Firefly activity was normalised to Renilla activity and is shown as percentages relative to wt FluPol_A/B_. The data shown are the mean ± SD of three independent experiments performed in technical duplicates. ***p ≤ 0.001 (two-way ANOVA; Dunnett’s multiple comparisons test).

CTD binding site 2A differs substantially between FluPol_B_ and FluPol_A_ (**Figure 1E** and **Figure 4A**). We introduced mutations at residues PA K289, R454 and S420, which are critical for CTD binding to FluPol_A_ and are not conserved in FluPol_B_, and we deleted the PA 550 loop,. which buttresses the CTD in FluPol_A_ and is considerably shortened in FluPol_B_. These modifications did not affect PA accumulation levels (**Figure 4B**) and specifically decreased *in vivo* CTD binding of FluPol_A_ but not FluPol_B_ (**Figure 4C** and **Figure S5B**). Consistently, the PA R454A and S420E mutations in the IBV background (IBV numbering: K450A and K416E) did not impair viral growth (**Figure 4D**) nor did PA K450A affect FluPol_B_ polymerase activity (**Figure 4E**). The PA S420E and PA 550 loop deletion impaired FluPol_B_ activity (**Figure 4E**), indicating that they hinder a function of the polymerase besides CTD binding.

**Figure 4.**
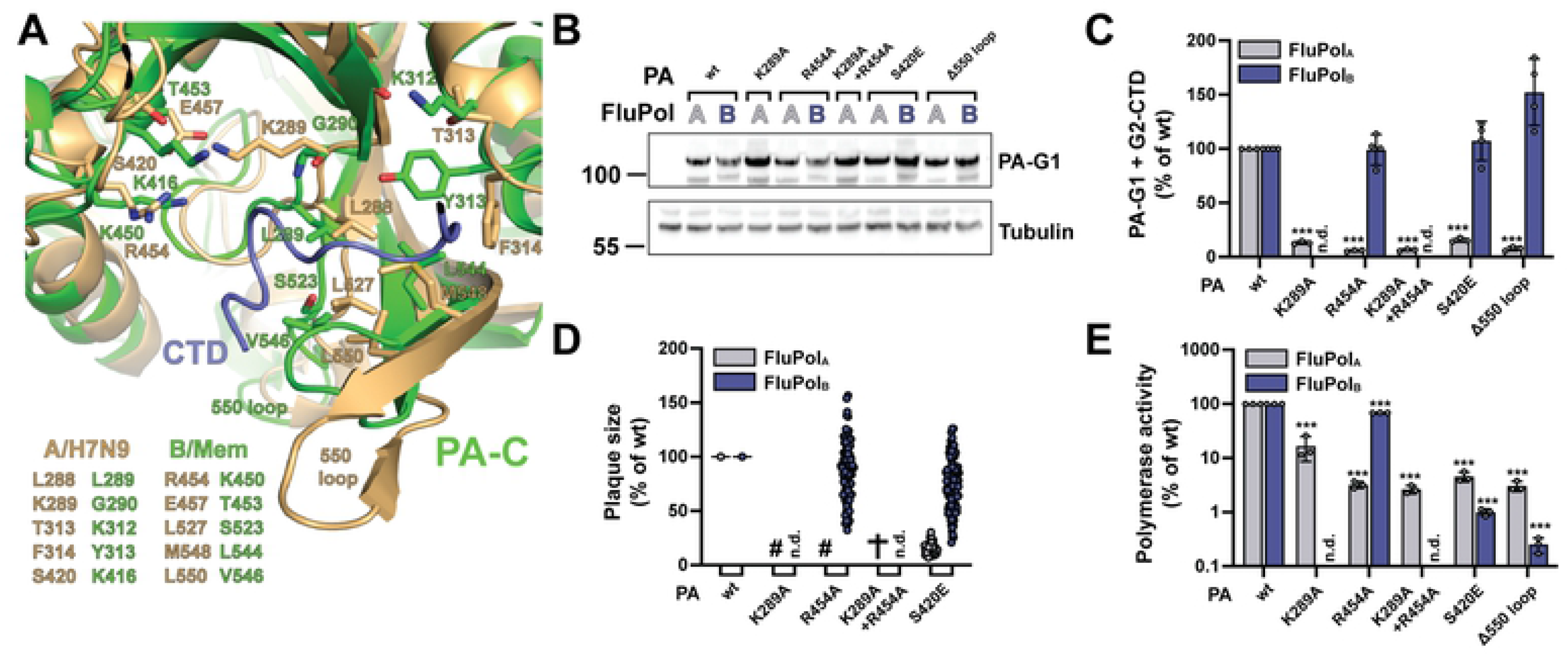
FluPol_B_ and FluPol_A_ CTD-binding mode at site 2A. **A.** Superposition of CTD peptide (slate blue tube) bound at site 2A of the PA subunit of FluPol_A_ (A/Zhejiang/DTID-ZJU01/2013(H7N9), green) with the equivalent region of FluPol_B_ (B/Memphis/13/2003, wheat), showing similarities and differences in CTD interacting residues. See sequence alignment Figure 1E. **B-E.** Protein expression (B), *in vivo* CTD binding (C), characterisation of recombinant IAV and IBV viruses (D) and polymerase activity (E) of CTD-binding site 2A mutants. Experiments were performed as described in Figure 3 for FluPol_B_ and FluPol_A_ site 1AB mutations. C. ***p ≤ 0.001 (two-way ANOVA; Dunnett’s multiple comparisons test). D. (#) not measurable pinhead-sized plaque diameter; (✞) no viral rescue, (n.d.) not determined. E. ***p ≤ 0.001 (two-way ANOVA; Dunnett’s multiple comparisons test).

### The PB2 627 domain is involved in CTD binding for FluPol_B_ but not FluPol_A_

The key CTD binding residues and the 3D structure of site 2B are often conserved between FluPol_B_ and FluPol_A_ (**Figure 1E** and **Figure 5A**). However, CTD binding at site 2B has never been observed *in vitro* with FluPol_A_, and the inserted PA 608 loop (IBV numbering) which buttresses the CTD at the junction between the PA-Cter and PB2 627 domains in FluPol_B_ is absent in FluPol_A_. The PA R608A mutation significantly decreased FluPol_B_ CTD binding in our cell-based complementation assay (**Figure 5B middle panel,** no counterpart residue in FluPol_A_). Consistently, the corresponding recombinant mutant IBV could not be rescued upon reverse genetics **(Figure S6)**, and the PA R608 mutant FluPol_B_ showed reduced polymerase activity (**Figure 5B, right panel**). We then mutated to alanines the residues W552 and R555 (W553 and R556 according to IBV numbering), which are located on the PB2 627 domain, make contact with the CTD pS5 in the FluPol_B_ co-crystal and are conserved between IBVs and IAVs (**Figure 1E** and **Figure 5A)**. The mutations did not affect PB2 accumulation levels (**Figure 5C**) and either decreased (R555A) or increased (W552A) CTD binding of FluPol_B_, whereas they had no effect on FluPol_A_ CTD binding (**Figure 5D** and **Figure S5C**). To rule out any CTD binding activity on the FluPol_A_ PB2 627 domain, we deleted the whole domain as described before [63]. The deletion strongly and specifically decreased CTD binding to FluPol_B_ but not to FluPol_A_ (**Figure 5E**). Nevertheless, single amino acid substitutions at residues PB2 W552 and R555 impaired viral growth and polymerase activity of FluPol_B_ as well as FluPol_A_, however with weaker effects on FluPol_A_ (**Figure 5F** and **5G**). Given the multiple functions attributed to the PB2 627 domain [63], the most likely interpretation of our data is that residues on the PB2 627 domain contribute to the CTD recruitment exclusively for IBVs while they have overlapping CTD-unrelated functions for IBVs and IAVs.

**Figure 5.**
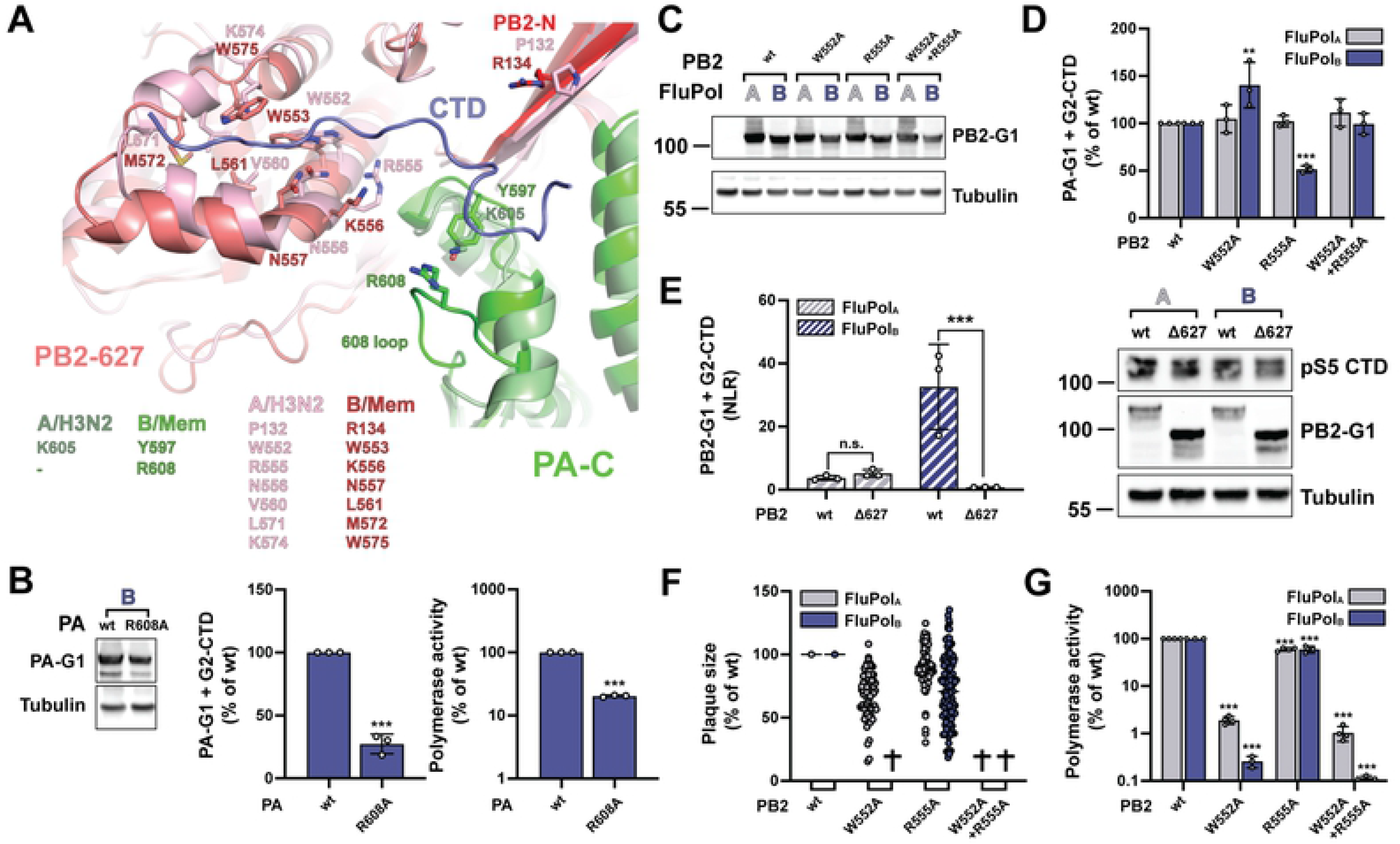
FluPol_A_ and FluPol_B_ CTD-binding mode at site 2B. **A.** Superposition of CTD peptide (slate blue tube) bound in site 2B of FluPol_B_ (B/Memphis/13/2003, PA green, PB2-N red, PB2-627 deep salmon) with the equivalent region of FluPol_A_ (A/NT/60/1968 (H3N2), [59], PDB: 6RR7, PA light green, PB2 pink), showing similarities and differences in CTD interacting residues. See sequence alignment Figure 1E. **B.** Protein expression (left), *in vivo* CTD binding (middle), and polymerase activity of FluPol_B_ PA R608A (right). Experiments were performed as described in Figure 3. The data shown are mean ± SD of three independent experiments performed in technical triplicates. ***p ≤ 0.001 (unpaired t test). **C-D.** Protein expression (C) and *in vivo* CTD binding (D) of CTD-binding site 2B mutants. Experiments were performed as described in Figure 3 for FluPol_B_ and FluPol_A_ site 1AB mutations. D. **p ≤ 0.002, ***p ≤ 0.001 (two-way ANOVA; Dunnett’s multiple comparisons test). **E.** *In vivo* CTD binding of PB2 Δ627 domain deletion mutants was investigated. G2-CTD was expressed by transient transfection in HEK-293T cells together with PB2-G1, PB1 and PA of FluPol_A_ (A/WSN/33, grey bars) or FluPol_B_ (B/Memphis/13/2003, blue bars). Luciferase activities were measured in cell lysates at 24 hpt. Normalised luciferase ratios (NLRs) were calculated as described in the Materials and Methods section. ***p ≤ 0.001 (two-way ANOVA; Sidak’s multiple comparisons test). Cell lysates were analysed in parallel by western blot with antibodies specific for the pS5 CTD, *G. princeps* luciferase (PB2-G1) and tubulin. **F-G.** Characterisation of recombinant IAV and IBV viruses (E) and polymerase activity (F) of CTD-binding site 2B mutants. Experiments were performed as described in Figure 3 for FluPol_B_ and FluPol_A_ site 1AB mutations. F. (✞) no viral rescue. G. ***p ≤ 0.001 (two-way ANOVA; Dunnett’s multiple comparisons test).

We asked whether this major difference between FluPol_B_ and FluPol_A_ CTD binding modes results in different levels of transcriptional activation by CTD mimicking peptides *in vitro*. A model has been proposed for FluPol_C_ in which the CTD stabilizes a transcription-competent conformation by binding at the interface of PB1, P3 (PA equivalent), and the flexible PB2 C-terminus [33]. Our observations suggest that the same model could apply to FluPol_B_ and not to FluPol_A_. Therefore, we tested the impact of pS5 CTD mimicking peptides of varying lengths (two, four, or six YSPTSPS repeats) on FluPol_B_ and FluPol_A_ *in vitro* transcriptional activity (**Figure 6A**). The FluPol_B_ *in vitro* endonuclease activity **(Figure 6A, lane 4)** and elongation activity **(Figure 6A, lane 8)** were increased in the presence of the six-repeat pS5 CTD mimicking peptide, and a similar trend was observed with FluPol_A_ **(Figure 6B, lanes 4 and 8)**. These data complement previous reports that CTD pS5 binding facilitates FluPol_A_ and FluPol_C_ transcriptional activity [33] and strengthen the hypothesis that the CTD stabilises FluPol in a transcription-competent conformation [20]. However, our finding that the FluPol_A_ PB2 627 domain has no CTD binding activity **(Figure 5G)** indicates that FluPol_A_ has evolved a divergent mechanism by which the CTD stabilizes the FluPol transcriptase. It also questions whether bridging of the PB2 627 and PA-Cter domains *per se* is needed for transcriptional activation of FluPols.

**Figure 6.**
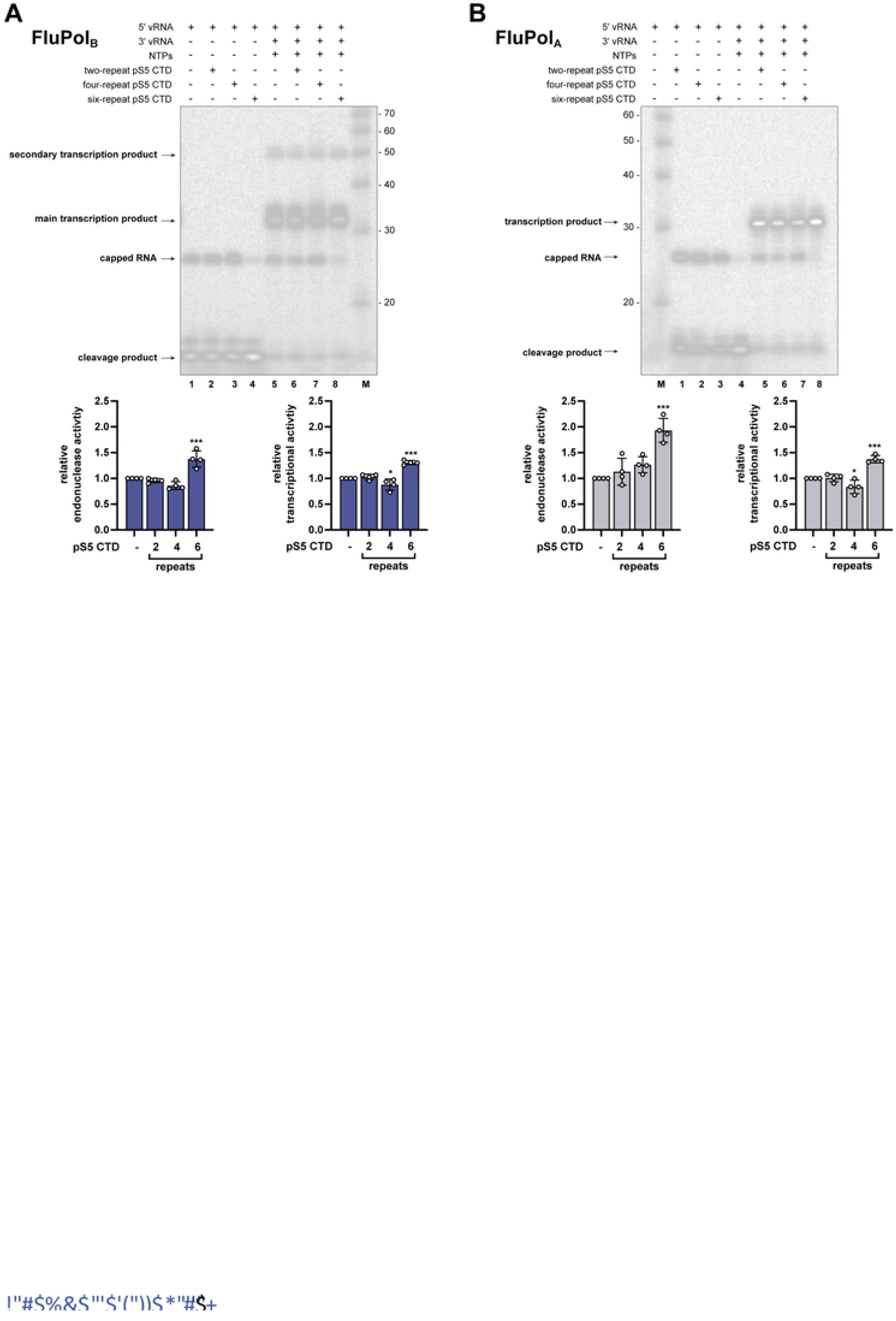
Effect of CTD pS5 peptides on *in vitro* endonuclease and transcription activity of FluPol_B_ and FluPol_A._ **A-B.** CTD pS5 peptides of different lengths (two, four and six repeats) were added to (A) FluPol_B_ (B/Memphis/13/2003) or (B) FluPol_A_ (bat influenza A (A/little yellow-shouldered bat/Guatemala/060/2010(H17N10)) *in vitro* activity reactions as described in Material and Methods. The left four lanes show endonuclease and right four lanes transcription reactions. Quantification of the reaction products of four independent experiments is shown below (FluPol_B_ in blue and FluPol_A_ in grey, respectively). The products of the reactions are normalised to the total RNA amount for each reaction and are presented as fractions of the activity of the reaction without peptide. *p ≤ 0.033, ***p ≤ 0.001 (one-way ANOVA; Dunnett’s multiple comparisons test).

### FluPol_B_ and FluPol_A_ bind to the host RNAP II independently of the CTD

The RNAP II transcriptional machinery is highly conserved across eukaryotes, and the CTD in particular shows almost no sequence differences among vertebrate (mammalian and avian) host species susceptible to IAV or IBV infection (**Figure S7**). It is therefore unlikely that differences in the CTD amino acid sequence drove the evolution of divergent IBV and IAV CTD-binding modes. We investigated whether FluPol_A/B_ can interact with the two major RNAP II subunits (RPB1, RPB2) independently of the CTD, using the split-gaussia luciferase complementation assay. The G2-RPB1 and RPB2-G2 fusion proteins were co-expressed with G1-tagged FluPol (PA-G1) by transient transfection as described above. Both combinations resulted in robust and comparable interaction signals (**Figure 7A)**. Interestingly, in the presence of a truncated RPB1 deleted from the CTD (RPB1ΔCTD), a stable interaction signal with FluPol_A/B_ could still be measured. Mutations in site 1A/B which reduced CTD binding to background levels (**Figure 3C**) had only weak effects on RPB1 and RPB1ΔCTD binding (PA K631A R635 in **Figure 7B,** PA K635A R638 in **Figure 7C**). The same was observed with site 2A mutations (PA K450A in **Figure 7B,** PA R454A in **Figure 7C)** and site 2B mutations and (PB2 R555A **Figure 7B,** PB2 K556A in **Figure 7C**). These findings, taken together with the relatively low affinity of FluPol for pS5 CTD peptides [32], suggest that the CTD is not the only interface between FluPol and the host RNAP II, and it may not be essential to connect FluPol to the RNAP II but rather to coordinate FluPol cap-snatching.

**Figure 7.**
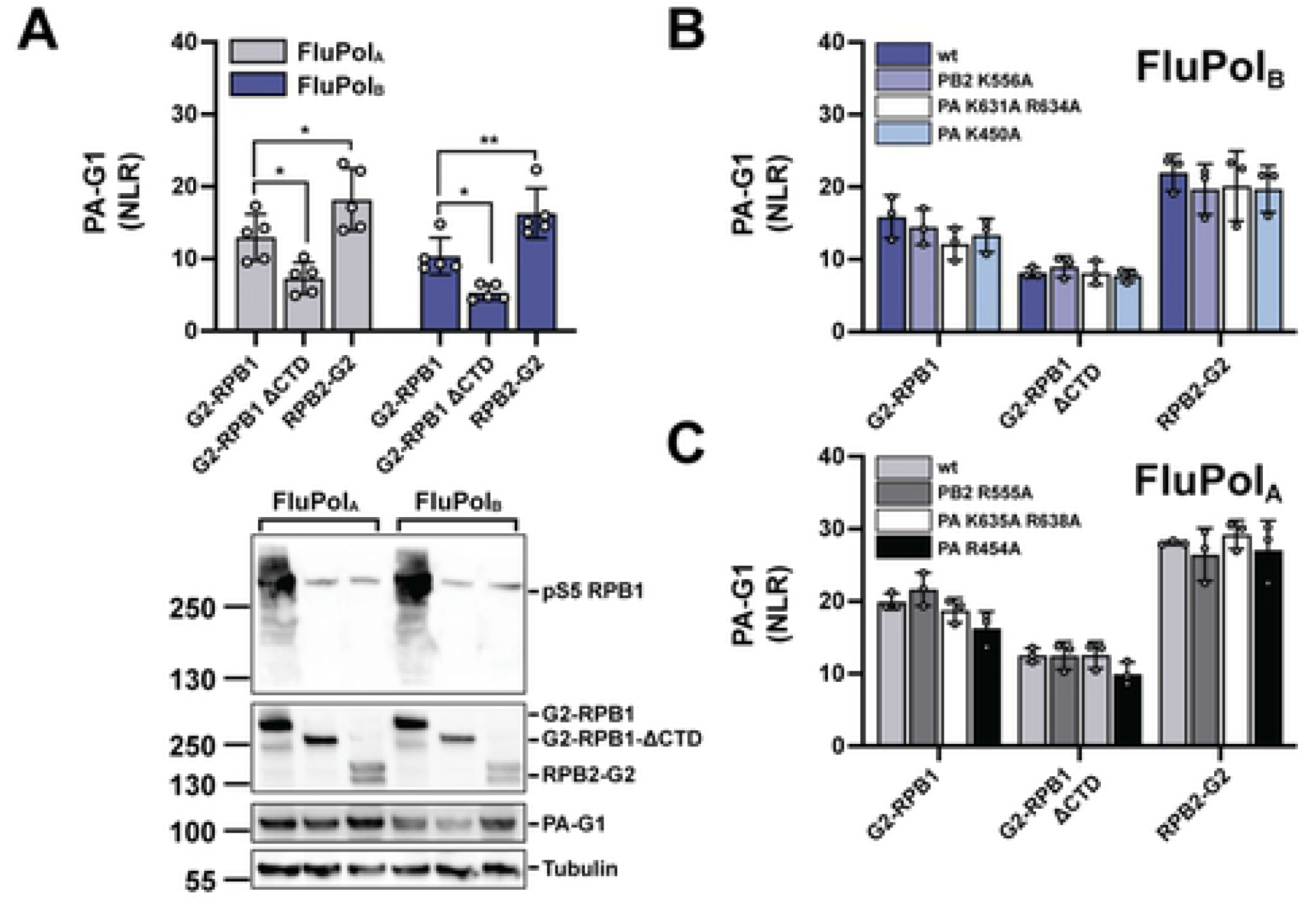
FluPol_B_ and FluPol_A_ binding to RPB1, RPB1ΔCTD and RPB2. **A.** Binding of FluPol_A_ (A/WSN/33, grey bars) and FluPol_B_ (B/Memphis/13/2003, blue bars) to RPB1, RPB1ΔCTD and RPB2 was evaluated. RPB1, RPB1ΔCTD and RPB2 were tagged to G2 and expressed by transient transfection in HEK-293T cells together with PB2, PB1 and PA-G1. Normalised luciferase activities (NLRs) were calculated as described in the Materials and Methods section. The data shown are the mean ± SD of five independent experiments performed in technical triplicates. *p ≤ 0.033, **p ≤ 0.002 (two-way ANOVA; Dunnett’s multiple comparisons test). Cell lysates were analysed in parallel by western blot with antibodies specific for the pS5 CTD, *G. princeps* luciferase and tubulin. **B-C.** Binding of (B) FluPol_B_ (B/Memphis/13/2003) and (C) FluPol_A_ (A/WSN/33) mutants in site 2B (PB2 K556/R555A), site 1 (PA K631A R634A/ PA K635A R638A) and site 2A (PA K450A / PA R454A) to RPB1, RPB1ΔCTD and RPB2 was evaluated as described in (A).

## Discussion

Here we report co-crystal structures of a human FluPol_B_ and an avian (isolated from human) FluPol_A_ bound to pS5 CTD mimicking peptides. We uncover the conformation and directionality of the CTD peptide bound to FluPol_B_ at a site that crosses over from the PA-Cter to the PB2 627 domain (site 2B), and has no counterpart on FluPol_A_ or FluPol_C_. Two CTD binding sites have been characterised on FluPol_A_ (sites 1A and 2A) ([32] and this study) and on FluPol_C_ (sites 1C and 2C, distinct from sites 1A and 2A) [33]. On the FluPol_B_ co-crystal structure, site 1B is similar to site 1A, whereas site 2B is distinct from site 2A and 2C.

By performing structure-based mutagenesis of FluPol_B_ and FluPol_A_ followed by a systematic investigation of FluPol-CTD binding, FluPol transcription/replication activity and viral phenotype, we confirm that CTD binding involves the same key residues at site 1AB for FluPol_B_ and FluPol_A_, but distinct and specific residues at site 2A for FluPol_A_ and site 2B for FluPol_B_, respectively. In particular, we demonstrate that the PA 606-609 loop, which buttresses the CTD at the junction between PA and PB2 in the FluPol_B_ co-crystal structure and is not conserved in FluPol_A_ or FluPol_C_, is an essential component of site 2B.

Our data and others’ [32, 33] demonstrate that IAVs, IBVs and ICVs have evolved divergent CTD binding modes, and raise questions about the driving force behind this divergent evolution. Large-scale meta-transcriptomic approaches have identified IBV-like and IDV-like viruses in fish and amphibians, suggesting that the influenza viruses of all four genera might be distributed among a much wider range of vertebrate species than recognised so far [57, 58]. Phylogenetic analyses, although limited by strong sampling biases across species, indicate that both virus-host co-divergence over long timescales and cross-species transmissions have shaped the evolution of influenza viruses. With one of the two CTD binding sites being conserved between IAVs and IBVs but absent in ICVs, the divergence of the bipartite CTD binding mode apparently matches the evolutionary distance between the three types of influenza viruses [64].

Interestingly however, we demonstrate that, in contrast to what is observed for IBVs and ICVs ([32, 33] and this study), the PB2 627 domain is not involved in CTD binding for IAVs. Therefore, from a mechanistic point of view, the CTD-dependent transcriptional activation of FluPol might be closer between IBVs and ICVs than between IBVs and IAVs as a consequence of a distinctive evolutionary pressure exerted on IAVs. The FluPol_A_ CTD binding mode presumably reflects an avian-optimised mode and co-evolved with protein interfaces between avian host factors and the PB2 627 domain, known to restrict avian IAV replication in humans (the principal hosts of IBVs and ICVs).

Another example of a functional interaction with RNAP II being achieved through distinct CTD binding is provided by the cellular mRNA capping enzyme (CE). The CEs from *Schizosaccharomyces pombe*, *Candida albicans* and *Mus musculus* were shown to bind directly S5 CTD repeats with very distinct binding interfaces and distinct conformations of the bound CTD [65]. These distantly related species show major differences in the CTD length and sequence [26] which could at least partially account for the divergence in CE-CTD binding modes. In contrast, the CTD is highly conserved among host species susceptible to IAV, IBV and ICV infections (**Figure S7**).

There is considerable evidence, however, that the FluPol-CTD interaction is only part of a more complex interaction pattern between the viral and cellular transcription machineries, raising the possibility that interactions between the FluPol and less conserved components of the cellular transcriptional machinery could have indirectly shaped the evolution of distinct CTD binding modes. We observed that a truncated RPB1 subunit, which lacks the CTD, retains partial binding to FluPol (**Figure 7**). Mass-spectrometry screenings have identified other RNAP II subunits and multiple transcriptional pausing and elongation factors as potential FluPol interaction partners [31]. Host factors involved in transcription such as DDX17 were found to bind FluPol and to determine IAV host-specificity [66]. By analogy, CEs not only bind to the pS5 CTD but also to the transcription pausing DRB Sensitivity-Inducing Factor (DSIF) [65] and make additional direct interactions with the nascent transcript exit site on the body of RNAP II [67]. Likewise, it was shown recently that the integrator complex binds RNAP II in its promotor-proximal paused state through direct interactions with the CTD of RPB1 but also with RPB2, RPB3, and RPB11, and through indirect interaction with the negative elongation factor NELF and DSIF [68]. Intriguingly, FluPol was also found to interact with the DSIF subunit SPT5 [69]. To what extent host-specific features of SPT5 or other cellular factors may have constrained the evolution of CTD-binding sites on FluPol remains to be explored.

We show that the *in vitro* transcriptional activity of FluPol_B_ is facilitated by the addition of CTD pS5 mimicking peptides, as reported previously for FluPol_A_ and FluPol_C_ [33]. The mechanism previously proposed for FluPol_C_ [20, 33] in which the CTD stabilises FluPol in a transcription-competent conformation by bridging P3 (the PA equivalent for ICVs) and PB2, could possibly apply to FluPol_B_ with PA-PB2 bridging occurring at site 2B. Our data show that it does not apply to FluPol_A_, unless another yet unidentified domain of PB2, distinct from the PB2 627 domain, is involved.

As underlined by the different sensitivity of IAV and IBVs to cap-binding inhibitors related to differences in the cap-binding mode of their PB2 subunits [70], a detailed understanding of structural and functional differences between FluPol_A_ and FluPol_B_ is of significant importance with regard to the development of broad-spectrum antivirals and need to be taken into account when targeting the FluPol-CTD binding interface for antiviral intervention.

## Acknowledgments

We thank Pr. Daniel Perez (University of Georgia), Dr. Bernard Delmas (INRAE) and Dr. Yves Janin (Institut Pasteur) for sharing material and reagents, and Dr. Yves Jacob for helpful discussions.

## Funding sources

This work was funded by the ANR grant FluTranscript (ANR-18-CE18-0028), holded jointly by SC and NN. TK was funded by the ANR grants Flutranscript ANR-18-CE18-0028 and Labex IBEID ANR-10-LABX-62-IBEID.

---

**Figure S1.**
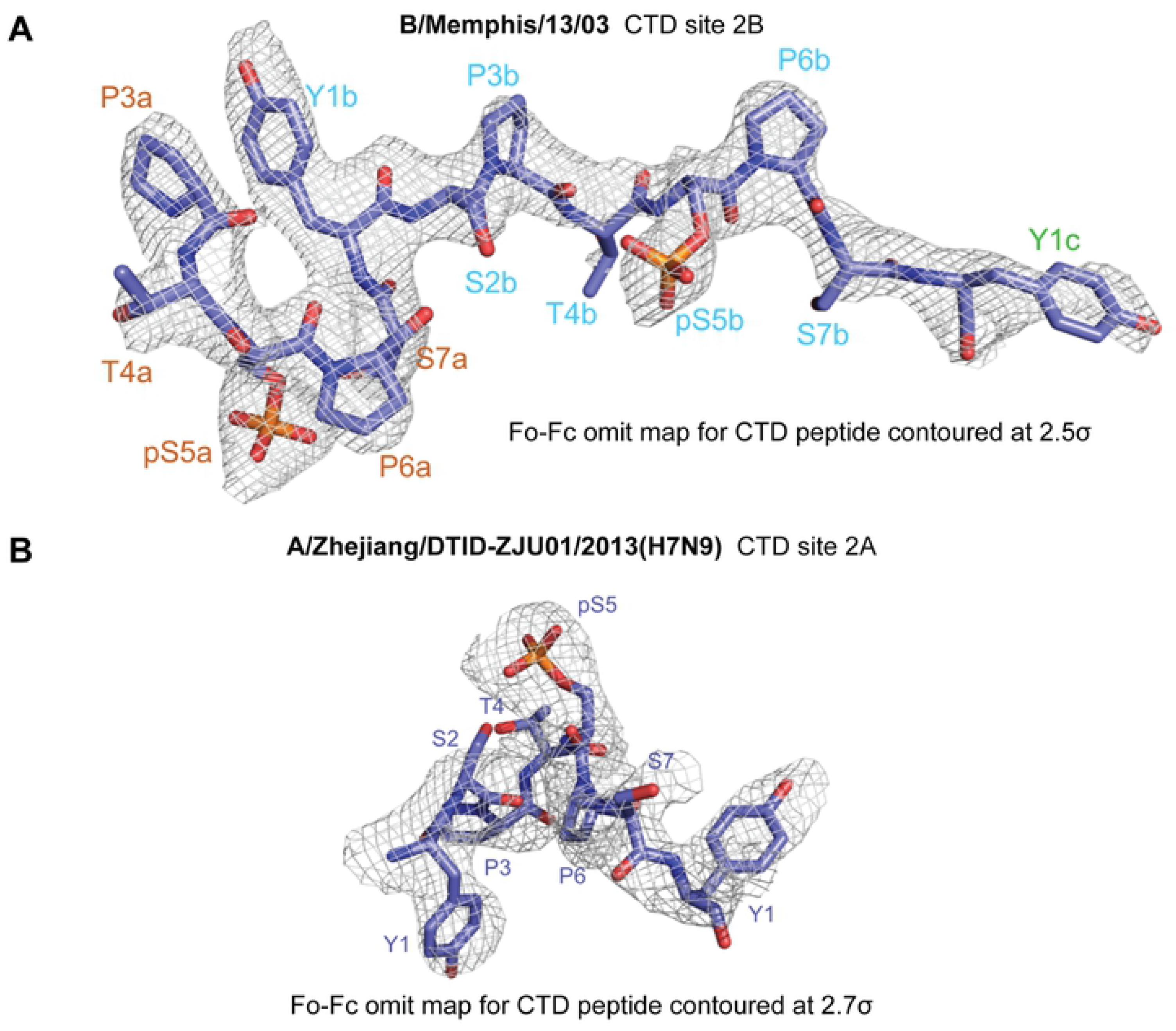
Omit maps for bound CTD peptide structures. **A.** CTD peptide bound in site 2B of FluPol_B_ (B/Memphis/13/2003) polymerase. Fo-Fc omit map shown at 2.5 σ with clear density for two phosphoserines (pS5a and pS5b). **B.** CTD peptide bound in site 2A of FluPol_A_ (A/Zhejiang/DTID-ZJU01/2013(H7N9)) polymerase. Fo-Fc omit map shown at 2.7 σ with clear density for one phosphoserine.

**Figure S2.**
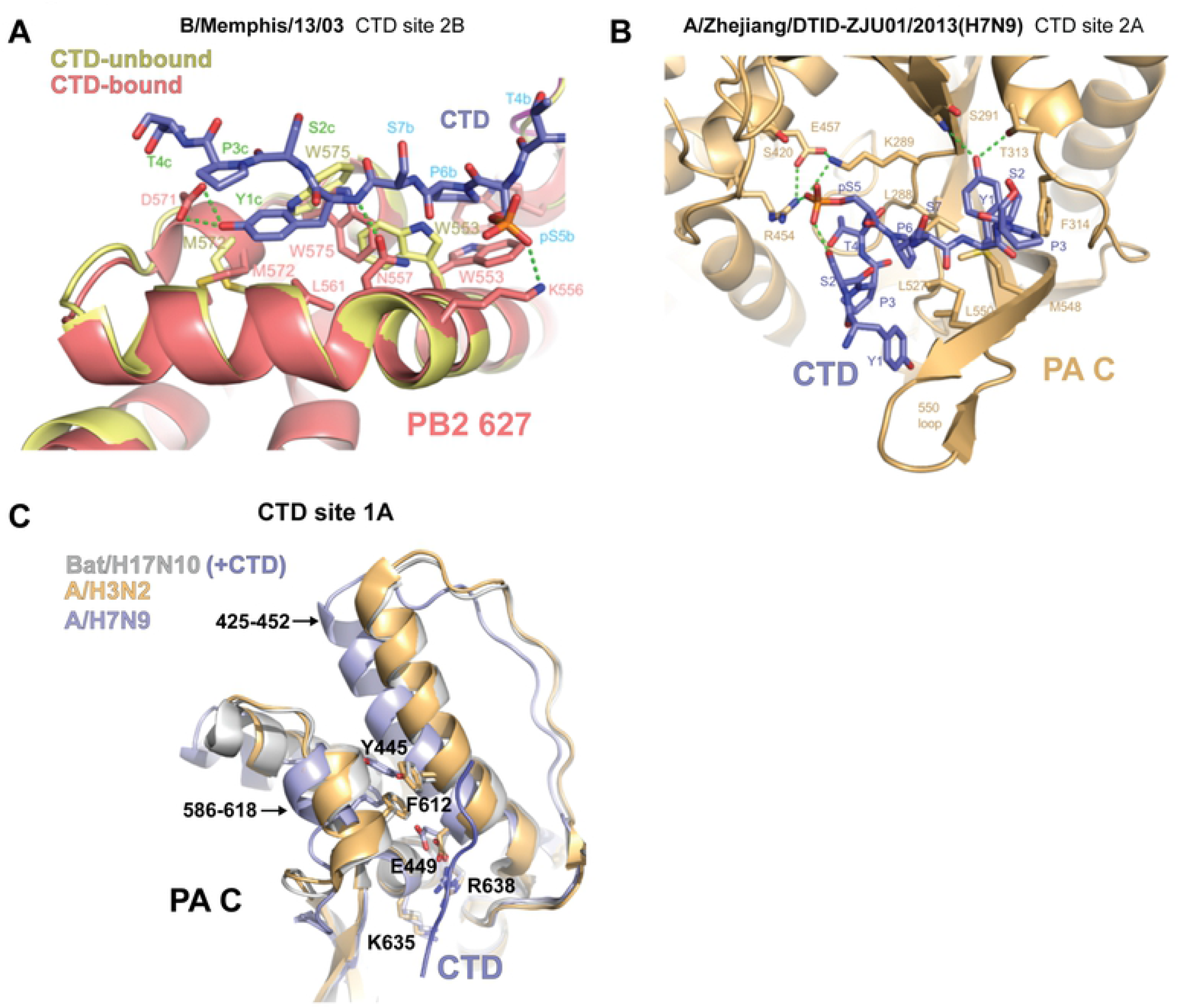
Structural analysis of CTD binding in FluPol_A_ and FluPol_B_. **A.** Superposition of the PB2-627 domains of CTD-bound (deep salmon with CTD in slate-blue) and unbound (wheat) FluPol_B_ (B/Memphis/13/2003) polymerase, showing induced-fit conformational changes of three key hydrophobic residues (PB2 W553, M572, W575). **B.** Details of the binding of the CTD peptide (slate blue) in site 2A of FluPol_A_ (A/Zhejiang/DTID-ZJU01/2013(H7N9) core) polymerase. **C.** Comparison of site 1A configuration for CTD bound form of FluPol_A_ (bat influenza A (A/little yellow-shouldered bat/Guatemala/060/2010(H17N10) [32], PDB: 5M3H, PA subunit light grey, CTD peptide slate-blue), CTD free, transcription active form of FluPol_A_ (A/NT/60/1968 (H3N2), [59], PDB: 6RR7, wheat) and dimeric FluPol_A_ (A/Zhejiang/DTID-ZJU01/2013(H7N9) core, light blue, this work). The FluPol_A_ (H7N9) polymerase core is the symmetrical dimer with each polymerase in the open, ‘dislocated’ state [18]. Due to the dislocation, PA regions 425-452 and 586-618 are rotated by ∼20°, which particularly effects the position of site 1A binding site residues Y445, E449 and F612. This likely explains the lack of CTD binding observed in site 1A for the H7N9 core, whereas site 2A is undistorted and occupied by CTD (Fig. S2B).

**Figure S3.**
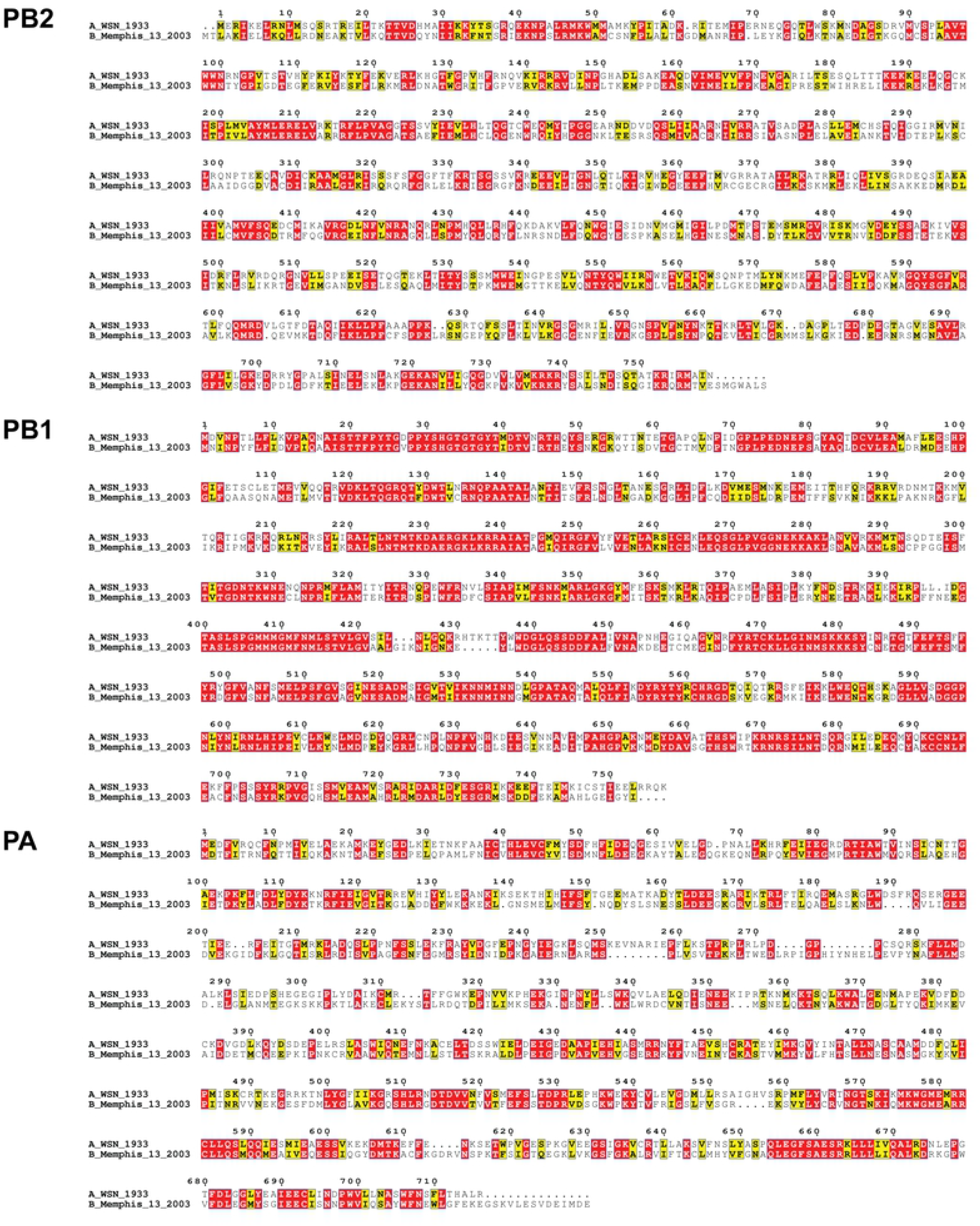
Sequence alignment of full-length PB2, PB1 and PA sequences of A/WSN/33 (A0A2Z5U3X0) and B/Memphis/13/2003 (Q5V8X3). Protein sequences were obtained from UniProt (https://www.uniprot.org/), aligned with SnapGene ® 6.0 and visualized by Espript 3.0 [55].

**Figure S4.**
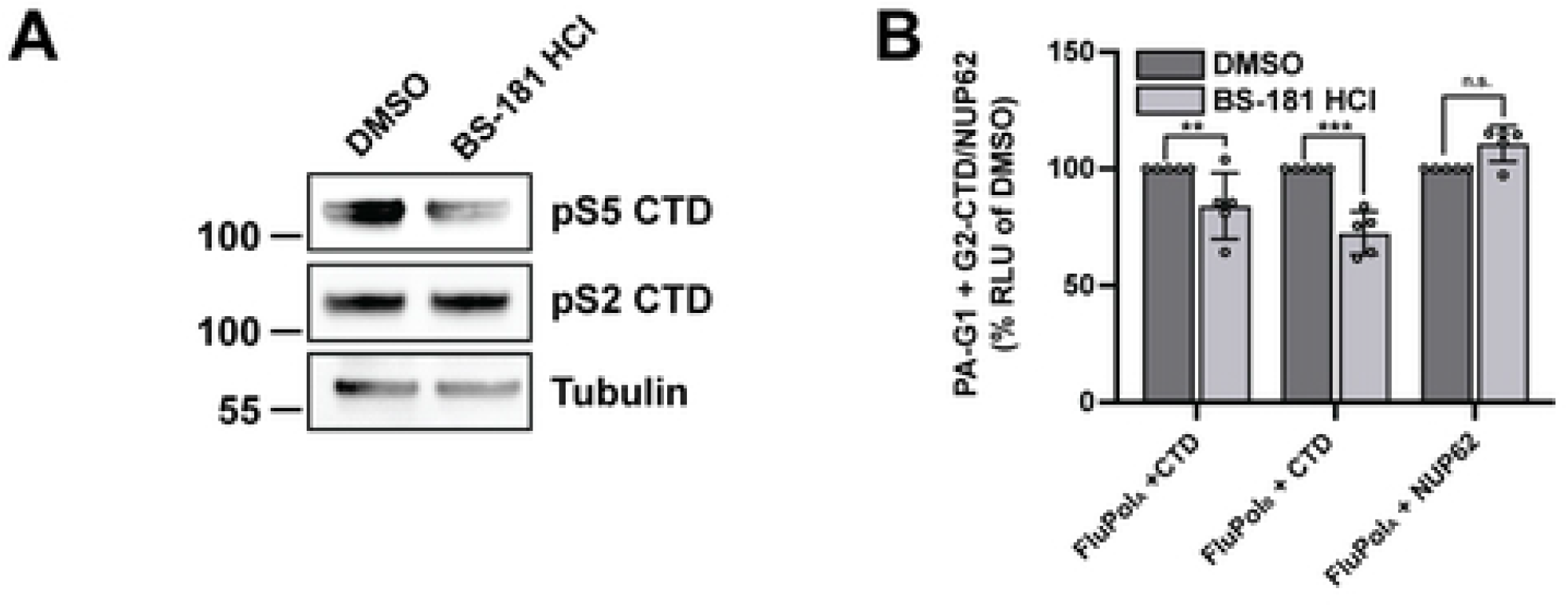
FluPol-CTD interaction in the presence of a CDK7 inhibitor. **A.** HEK-293T cells were transfected with G2-CTD. At 24 hpt cells were treated for 1 h with 20 µM BS-181-HCl (DMSO final concentration 0.2 %). Cell lysates were analysed by western blot with antibodies specific for pS5 or pS2 CTD and tubulin. **_B._** *In vivo* CTD binding of FluPol_A_ (A/WSN/33) and FluPol_B_ (B/Memphis/13/2003). G2-tagged CTD was expressed by transient transfection in HEK-293T cells together with the viral polymerase subunit PB2, PB1 and PA-G1. At 24 hpt cells were treated for 1 h with 20 µM BS-181-HCl or 0.2 % DMSO before cell lysis and measurement of *G. princeps* luciferase activity as described in the Material and Methods section. As a control, the previously described FluPol_A_– NUP62 interaction was investigated by co-transfection of G2-NUP62, PB2, PB1 and PA-G1. RLUs are expressed as percentages relative to DMSO treated cells. The data shown are mean ± SD of five independent experiments performed in technical triplicates. ***p ≤ 0.002, ***p ≤ 0.001 (two-way ANOVA; Dunnett’s multiple comparisons test).

**Figure S5:**
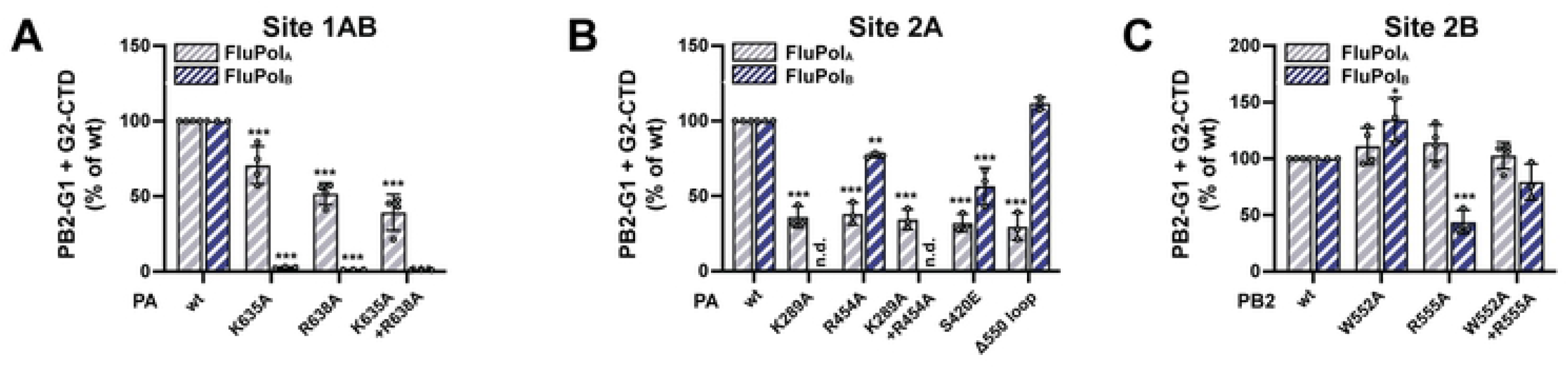
*In vivo* CTD binding of FluPol_A_ and FluPol_B_ mutants. **A-C.** *In vivo* CTD binding of the indicated site 1AB (A), site 2A (B) and site 2B (C) mutants of FluPol_A_ (A/WSN/33, grey hatched bars) and FluPol_B_ (B/Memphis/13/2003, blue hatched bars). The G2-tagged CTD was expressed by transient transfection in HEK-293T cells together with PB2-G1, PB1 and PA. RLUs are expressed as percentages relative to wt FluPol_A/B_. The data shown are the mean ± SD of at least three independent experiments performed in technical triplicates. ***p ≤ 0.002, ***p ≤ 0.001 (two-way ANOVA; Dunnett’s multiple comparisons test). (n.d.) not determined.

**Figure S6:**
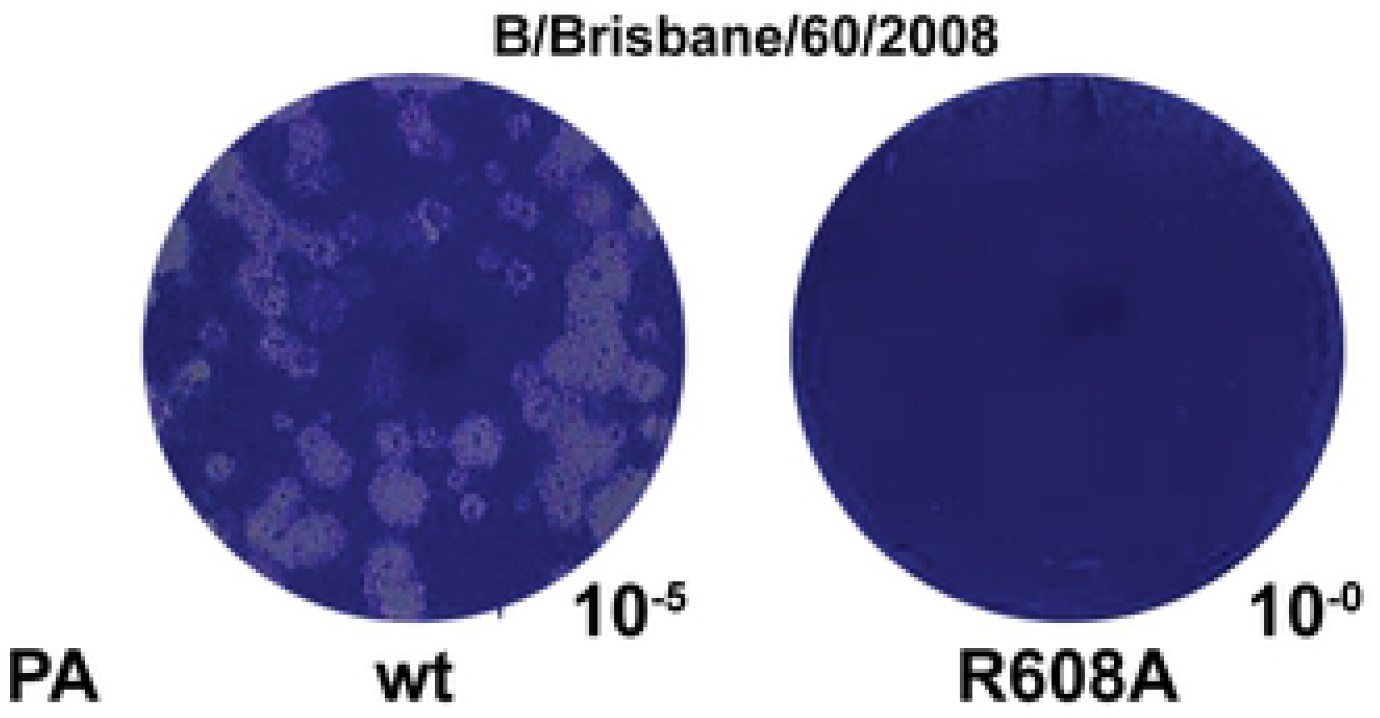
Plaque phenotype of FluPolB PA R608A. Characterisation of recombinant IBV (B/Brisbane/60/2008) PA R608A mutant virus. Recombinant viruses with the indicated mutations were generated by reverse genetics as described in the Material and Methods section. Reverse genetic supernatants were titrated on MDCK cells and stained at 72 hpi by crystal violet. The pictures show one representative plaque assay with the indicated ten-fold dilution.

**Figure S7:**
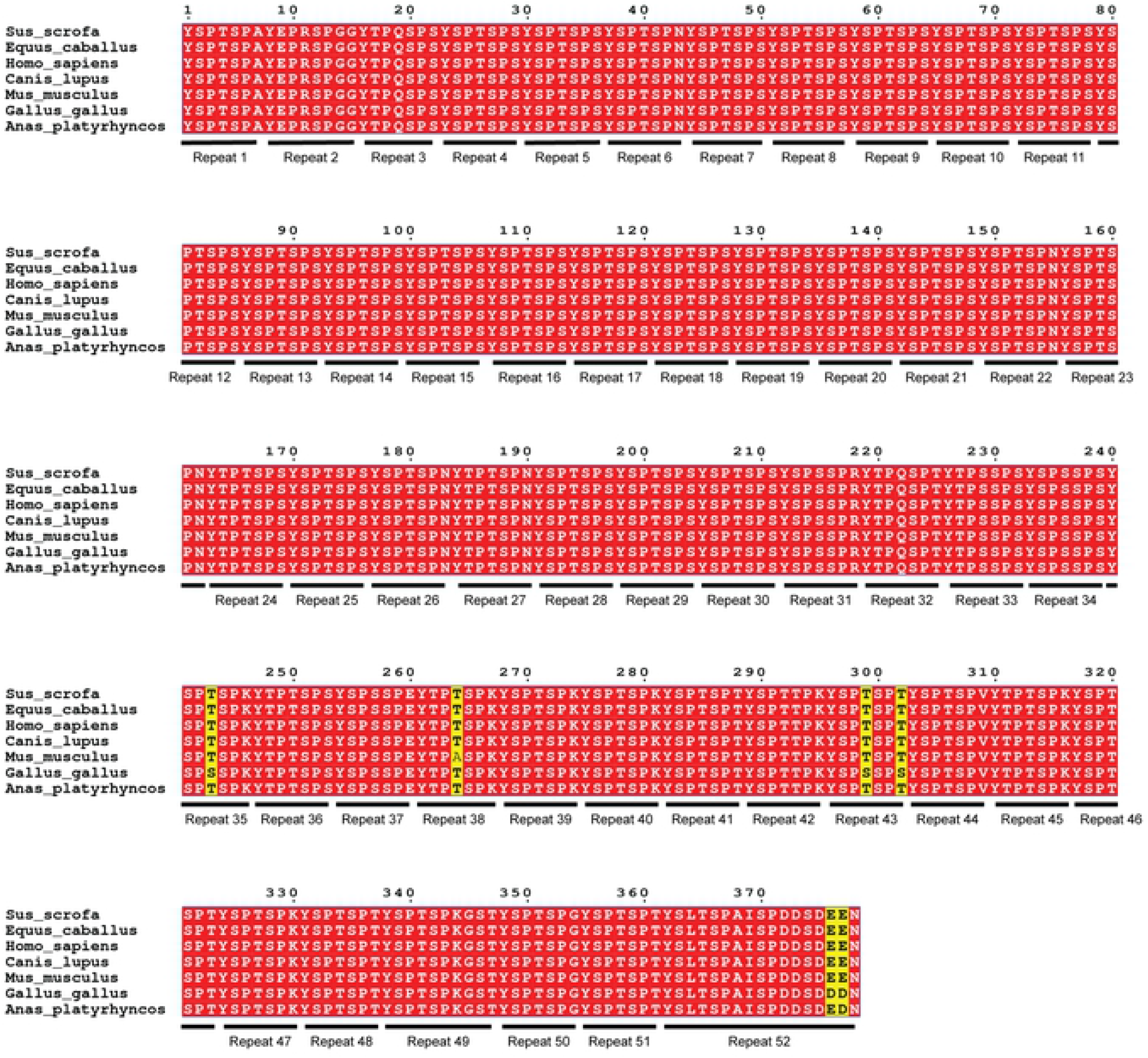
Sequence alignment of the RPB1 C-terminal domain (CTD) across species. The RPB1 CTD sequences of *Sus scrofa* (wild boar), *Equus caballus* (horse), *Homo sapiens* (human*), Canis lupus* (wolf), *Mus musculus* (house mouse), *Gallus gallus* (chicken) and *Anas platyrhynchos* (wild duck) were obtained as described in the Material and Methods section, aligned with SnapGene ® 6.0 and visualised by Espript 3.0 [55]. The CTD repeat numbers are indicated below the sequence alignment. Identical and similar residues are indicated in red or yellow, respectively.

## Notes

### Competing Interest Statement

The authors have declared no competing interest.

## References

1. Krammer F, Smith GJD, Fouchier RAM, Peiris M, Kedzierska K, Doherty PC, et al. Influenza. Nat Rev Dis Prim. 2018;4: 3. doi:10.1038/s41572-018-0002-y

2. Caini S, Kroneman M, Wiegers T, El Guerche-Séblain C, Paget J. Clinical characteristics and severity of influenza infections by virus type, subtype, and lineage: A systematic literature review. Influenza Other Respi Viruses. 2018;12: 780–792. doi:10.1111/IRV.12575

3. Zaraket H, Hurt AC, Clinch B, Barr I, Lee N. Burden of influenza B virus infection and considerations for clinical management. Antiviral Res. 2021;185. doi:10.1016/J.ANTIVIRAL.2020.104970

4. Yoon SW, Webby RJ, Webster RG. Evolution and ecology of influenza A viruses. Curr Top Microbiol Immunol. 2014;385: 359–375. doi:10.1007/82_2014_396

5. Suzuki Y, Nei M. Origin and evolution of influenza virus hemagglutinin genes. Mol Biol Evol. 2002;19: 501–509. doi:10.1093/OXFORDJOURNALS.MOLBEV.A004105

6. Parry R, Wille M, Turnbull OMH, Geoghegan JL, Holmes EC. Divergent Influenza-Like Viruses of Amphibians and Fish Support an Ancient Evolutionary Association. Viruses. 2020;12. doi:10.3390/V12091042

7. Baker SF, Nogales A, Finch C, Tuffy KM, Domm W, Perez DR, et al. Influenza A and B virus intertypic reassortment through compatible viral packaging signals. J Virol. 2014;88: 10778– 10791. doi:10.1128/JVI.01440-14

8. Eisfeld AJ, Neumann G, Kawaoka Y. At the centre: influenza A virus ribonucleoproteins. Nat Rev Microbiol. 2015;13: 28–41. doi:10.1038/nrmicro3367

9. Coloma R, Arranz R, de la Rosa-Trevín JM, Sorzano COS, Munier S, Carlero D, et al. Structural insights into influenza A virus ribonucleoproteins reveal a processive helical track as transcription mechanism. Nat Microbiol. 2020;5: 727–734. doi:10.1038/s41564-020-0675-3

10. Moeller A, Kirchdoerfer RN, Potter CS, Carragher B, Wilson IA. Organization of the influenza virus replication machinery. Science. 2012;338: 1631–1634. doi:10.1126/SCIENCE.1227270

11. Arranz R, Coloma R, Chichón FJ, Conesa JJ, Carrascosa JL, Valpuesta JM, et al. The structure of native influenza virion ribonucleoproteins. Science. 2012;338: 1634–1637. doi:10.1126/SCIENCE.1228172

12. Wandzik JM, Kouba T, Cusack S. Structure and Function of Influenza Polymerase. Cold Spring Harb Perspect Med. 2021;11. doi:10.1101/CSHPERSPECT.A038372

13. Deng T, Vreede FT, Brownlee GG. Different de novo initiation strategies are used by influenza virus RNA polymerase on its cRNA and viral RNA promoters during viral RNA replication. J Virol. 2006;80: 2337–48. doi:10.1128/JVI.80.5.2337-2348.2006

14. Oymans J, Te Velthuis AJW. A Mechanism for Priming and Realignment during Influenza A Virus Replication. J Virol. 2018;92. doi:10.1128/JVI.01773-17

15. Plotch SJ, Bouloy M, Krug RM. Transfer of 5’-terminal cap of globin mRNA to influenza viral complementary RNA during transcription in vitro. Proc Natl Acad Sci U S A. 1979;76: 1618– 22. doi:10.1073/pnas.76.4.1618

16. Bouloy M, Plotch SJ, Krug RM. Both the 7-methyl and the 2’-O-methyl groups in the cap of mRNA strongly influence its ability to act as primer for influenza virus RNA transcription. Proc Natl Acad Sci U S A. 1980;77: 3952–6. doi:10.1073/pnas.77.7.3952

17. Poon LL, Pritlove DC, Fodor E, Brownlee GG. Direct evidence that the poly(A) tail of influenza A virus mRNA is synthesized by reiterative copying of a U track in the virion RNA template. J Virol. 1999;73: 3473–6. doi:10.1128/JVI.73.4.3473-3476.1999

18. Wandzik JM, Kouba T, Karuppasamy M, Pflug A, Drncova P, Provaznik J, et al. A Structure-Based Model for the Complete Transcription Cycle of Influenza Polymerase. Cell. 2020;181: 877–893.e21. doi:10.1016/j.cell.2020.03.061

19. Bouloy M, Plotch SJ, Krug RM. Globin mRNAs are primers for the transcription of influenza viral RNA in vitro. Proc Natl Acad Sci U S A. 1978;75: 4886–90. doi:10.1073/pnas.75.10.4886

20. Walker AP, Fodor E. Interplay between Influenza Virus and the Host RNA Polymerase II Transcriptional Machinery. Trends Microbiol. 2019;27: 398–407. doi:10.1016/j.tim.2018.12.013

21. Dias A, Bouvier D, Crépin T, McCarthy AA, Hart DJ, Baudin F, et al. The cap-snatching endonuclease of influenza virus polymerase resides in the PA subunit. Nature. 2009;458: 914– 8. doi:10.1038/nature07745

22. Kouba T, Drncová P, Cusack S. Structural snapshots of actively transcribing influenza polymerase. Nat Struct Mol Biol. 2019;26: 460–470. doi:10.1038/s41594-019-0232-z

23. Worch R, Niedzwiecka A, Stepinski J, Mazza C, Jankowska-Anyszka M, Darzynkiewicz E, et al. Specificity of recognition of mRNA 5’ cap by human nuclear cap-binding complex. RNA. 2005;11: 1355–63. doi:10.1261/rna.2850705

24. Cramer P, Bushnell DA, Kornberg RD. Structural basis of transcription: RNA polymerase II at 2.8 angstrom resolution. Science. 2001;292: 1863–1876. doi:10.1126/SCIENCE.1059493

25. Schüller R, Forné I, Straub T, Schreieck A, Texier Y, Shah N, et al. Heptad-Specific Phosphorylation of RNA Polymerase II CTD. Mol Cell. 2016;61: 305–14. doi:10.1016/j.molcel.2015.12.003

26. Eick D, Geyer M. The RNA polymerase II carboxy-terminal domain (CTD) code. Chem Rev. 2013;113: 8456–90. doi:10.1021/cr400071f

27. Martínez-Alonso M, Hengrung N, Fodor E. RNA-Free and Ribonucleoprotein-Associated Influenza Virus Polymerases Directly Bind the Serine-5-Phosphorylated Carboxyl-Terminal Domain of Host RNA Polymerase II. J Virol. 2016;90: 6014–6021. doi:10.1128/JVI.00494-16

28. Engelhardt OG, Smith M, Fodor E. Association of the influenza A virus RNA-dependent RNA polymerase with cellular RNA polymerase II. J Virol. 2005;79: 5812–8. doi:10.1128/JVI.79.9.5812-5818.2005

29. Vos SM, Farnung L, Urlaub H, Cramer P. Structure of paused transcription complex Pol II-DSIF-NELF. Nature. 2018;560: 601–606. doi:10.1038/s41586-018-0442-2

30. Chan AY, Vreede FT, Smith M, Engelhardt OG, Fodor E. Influenza virus inhibits RNA polymerase II elongation. Virology. 2006;351: 210–7. doi:10.1016/j.virol.2006.03.005

31. Krischuns T, Lukarska M, Naffakh N, Cusack S. Influenza Virus RNA-Dependent RNA Polymerase and the Host Transcriptional Apparatus. Annu Rev Biochem. 2021;90: 321–348. doi:10.1146/ANNUREV-BIOCHEM-072820-100645

32. Lukarska M, Fournier G, Pflug A, Resa-Infante P, Reich S, Naffakh N, et al. Structural basis of an essential interaction between influenza polymerase and Pol II CTD. Nature. 2017;541: 117– 121. doi:10.1038/nature20594

33. Serna Martin I, Hengrung N, Renner M, Sharps J, Martínez-Alonso M, Masiulis S, et al. A Mechanism for the Activation of the Influenza Virus Transcriptase. Mol Cell. 2018;70: 1101–1110.e4. doi:10.1016/j.molcel.2018.05.011

34. McCoy AJ, Grosse-Kunstleve RW, Adams PD, Winn MD, Storoni LC, Read RJ. Phaser crystallographic software. J Appl Crystallogr. 2007;40: 658–674. doi:10.1107/S0021889807021206

35. Thierry E, Guilligay D, Kosinski J, Bock T, Gaudon S, Round A, et al. Influenza Polymerase Can Adopt an Alternative Configuration Involving a Radical Repacking of PB2 Domains. Mol Cell. 2016;61: 125–37. doi:10.1016/j.molcel.2015.11.016

36. Emsley P, Cowtan K. Coot: model-building tools for molecular graphics. Acta Crystallogr D Biol Crystallogr. 2004;60: 2126–2132. doi:10.1107/S0907444904019158

37. Murshudov GN, Vagin AA, Dodson EJ. Refinement of macromolecular structures by the maximum-likelihood method. Acta Crystallogr D Biol Crystallogr. 1997;53: 240–255. doi:10.1107/S0907444996012255

38. Chen VB, Arendall WB, Headd JJ, Keedy DA, Immormino RM, Kapral GJ, et al. MolProbity: all-atom structure validation for macromolecular crystallography. Acta Crystallogr D Biol Crystallogr. 2010;66: 12–21. doi:10.1107/S0907444909042073

39. Wandzik JM, Kouba T, Cusack S. Structure and Function of Influenza Polymerase. Cold Spring Harb Perspect Med. 2020; a038372. doi:10.1101/cshperspect.a038372

40. Fodor E, Devenish L, Engelhardt OG, Palese P, Brownlee GG, García-Sastre A. Rescue of influenza A virus from recombinant DNA. J Virol. 1999;73: 9679–9682. doi:10.1128/JVI.73.11.9679-9682.1999

41. Nogales A, Perez DR, Santos J, Finch C, Martínez-Sobrido L. Reverse Genetics of Influenza B Viruses. Methods Mol Biol. 2017;1602: 205–238. doi:10.1007/978-1-4939-6964-7_14

42. Reich S, Guilligay D, Pflug A, Malet H, Berger I, Crépin T, et al. Structural insight into cap-snatching and RNA synthesis by influenza polymerase. Nature. 2014;516: 361–6. doi:10.1038/nature14009

43. Cassonnet P, Rolloy C, Neveu G, Vidalain PO, Chantier T, Pellet J, et al. Benchmarking a luciferase complementation assay for detecting protein complexes. Nat Methods. 2011;8: 990– 992. doi:10.1038/NMETH.1773

44. Munier S, Rolland T, Diot C, Jacob Y, Naffakh N. Exploration of Binary Virus–Host Interactions Using an Infectious Protein Complementation Assay. Mol Cell Proteomics. 2013;12: 2845. doi:10.1074/MCP.M113.028688

45. Zheng L, Baumann U, Reymond JL. An efficient one-step site-directed and site-saturation mutagenesis protocol. Nucleic Acids Res. 2004;32. doi:10.1093/NAR/GNH110

46. Krischuns T, Günl F, Henschel L, Binder M, Willemsen J, Schloer S, et al. Phosphorylation of TRIM28 Enhances the Expression of IFN-β and Proinflammatory Cytokines During HPAIV Infection of Human Lung Epithelial Cells. Front Immunol. 2018;9: 2229. doi:10.3389/fimmu.2018.02229

47. Matrosovich M, Matrosovich T, Garten W, Klenk HD. New low-viscosity overlay medium for viral plaque assays. Virol J. 2006;3. doi:10.1186/1743-422X-3-63

48. Schindelin J, Arganda-Carreras I, Frise E, Kaynig V, Longair M, Pietzsch T, et al. Fiji: an open-source platform for biological-image analysis. Nat Methods. 2012;9: 676–682. doi:10.1038/NMETH.2019

49. O’Leary NA, Wright MW, Brister JR, Ciufo S, Haddad D, McVeigh R, et al. Reference sequence (RefSeq) database at NCBI: current status, taxonomic expansion, and functional annotation. Nucleic Acids Res. 2016;44: D733–D745. doi:10.1093/NAR/GKV1189

50. Buchfink B, Reuter K, Drost HG. Sensitive protein alignments at tree-of-life scale using DIAMOND. Nat Methods. 2021;18: 366–368. doi:10.1038/S41592-021-01101-X

51. Grabherr MG, Haas BJ, Yassour M, Levin JZ, Thompson DA, Amit I, et al. Full-length transcriptome assembly from RNA-Seq data without a reference genome. Nat Biotechnol. 2011;29: 644–652. doi:10.1038/NBT.1883

52. Kim D, Paggi JM, Park C, Bennett C, Salzberg SL. Graph-based genome alignment and genotyping with HISAT2 and HISAT-genotype. Nat Biotechnol. 2019;37: 907–915. doi:10.1038/S41587-019-0201-4

53. Li H. Minimap2: pairwise alignment for nucleotide sequences. Bioinformatics. 2018;34: 3094–3100. doi:10.1093/BIOINFORMATICS/BTY191

54. Kovaka S, Zimin A V., Pertea GM, Razaghi R, Salzberg SL, Pertea M. Transcriptome assembly from long-read RNA-seq alignments with StringTie2. Genome Biol. 2019;20. doi:10.1186/S13059-019-1910-1

55. Robert X, Gouet P. Deciphering key features in protein structures with the new ENDscript server. Nucleic Acids Res. 2014;42. doi:10.1093/NAR/GKU316

56. Reich S, Guilligay D, Cusack S. An in vitro fluorescence based study of initiation of RNA synthesis by influenza B polymerase. Nucleic Acids Res. 2017;45: 3353–3368. doi:10.1093/NAR/GKX043

57. Shi M, Lin XD, Chen X, Tian JH, Chen LJ, Li K, et al. The evolutionary history of vertebrate RNA viruses. Nature. 2018;556: 197–202. doi:10.1038/S41586-018-0012-7

58. Parry R, Wille M, Turnbull OMH, Geoghegan JL, Holmes EC. Divergent Influenza-Like Viruses of Amphibians and Fish Support an Ancient Evolutionary Association. Viruses. 2020;12. doi:10.3390/V12091042

59. Fan H, Walker AP, Carrique L, Keown JR, Serna Martin I, Karia D, et al. Structures of influenza A virus RNA polymerase offer insight into viral genome replication. Nature. 2019;573: 287–290. doi:10.1038/s41586-019-1530-7

60. Chang S, Sun D, Liang H, Wang J, Li J, Guo L, et al. Cryo-EM structure of influenza virus RNA polymerase complex at 4.3 Å resolution. Mol Cell. 2015;57: 925–935. doi:10.1016/J.MOLCEL.2014.12.031

61. Biswas SK, Nayak DP. Mutational analysis of the conserved motifs of influenza A virus polymerase basic protein 1. J Virol. 1994;68: 1819–1826. doi:10.1128/JVI.68.3.1819-1826.1994

62. Ali S, Heathcote DA, Kroll SHB, Jogalekar AS, Scheiper B, Patel H, et al. The development of a selective cyclin-dependent kinase inhibitor that shows antitumor activity. Cancer Res. 2009;69: 6208–6215. doi:10.1158/0008-5472.CAN-09-0301

63. Nilsson BE, Velthuis AJW te, Fodor E. Role of the PB2 627 Domain in Influenza A Virus Polymerase Function. J Virol. 2017;91: 2467–2483. doi:10.1128/JVI.02467-16

64. Wille M, Holmes EC. The Ecology and Evolution of Influenza Viruses. Cold Spring Harb Perspect Med. 2020;10: 1–19. doi:10.1101/CSHPERSPECT.A038489

65. Doamekpor SK, Sanchez AM, Schwer B, Shuman S, Lima CD. How an mRNA capping enzyme reads distinct RNA polymerase II and Spt5 CTD phosphorylation codes. Genes Dev. 2014;28: 1323–36. doi:10.1101/gad.242768.114

66. Bortz E, Westera L, Maamary J, Steel J, Albrecht RA, Manicassamy B, et al. Host- and strain-specific regulation of influenza virus polymerase activity by interacting cellular proteins. MBio. 2011;2: 1–10. doi:10.1128/mBio.00151-11

67. Martinez-Rucobo FW, Kohler R, van de Waterbeemd M, Heck AJR, Hemann M, Herzog F, et al. Molecular Basis of Transcription-Coupled Pre-mRNA Capping. Mol Cell. 2015;58: 1079– 89. doi:10.1016/j.molcel.2015.04.004

68. Fianu I, Chen Y, Dienemann C, Dybkov O, Linden A, Urlaub H, et al. Structural basis of Integrator-mediated transcription regulation. Science. 2021;374: 883–887. doi:10.1126/SCIENCE.ABK0154

69. Bradel-Tretheway BG, Mattiacio JL, Krasnoselsky A, Stevenson C, Purdy D, Dewhurst S, et al. Comprehensive proteomic analysis of influenza virus polymerase complex reveals a novel association with mitochondrial proteins and RNA polymerase accessory factors. J Virol. 2011;85: 8569–81. doi:10.1128/JVI.00496-11

70. Pflug A, Gaudon S, Resa-Infante P, Lethier M, Reich S, Schulze WM, et al. Capped RNA primer binding to influenza polymerase and implications for the mechanism of cap-binding inhibitors. Nucleic Acids Res. 2018;46: 956–971. doi:10.1093/NAR/GKX1210

